# Simple and efficient differentiation of human iPSCs into contractible skeletal muscles for muscular disease modeling

**DOI:** 10.1101/2021.11.22.468571

**Authors:** Muhammad Irfanur Rashid, Takuji Ito, Daisuke Shimojo, Kanae Arimoto, Kazunari Onodera, Rina Okada, Takunori Nagashima, Kazuki Yamamoto, Zohora Khatun, Hideyuki Okano, Hidetoshi Sakurai, Kazunori Shimizu, Manabu Doyu, Yohei Okada

## Abstract

Pathophysiological analysis and drug discovery targeting human diseases require disease models that suitably recapitulate patients’ pathology. Disease-specific human induced pluripotent stem cells (hiPSCs) can potentially recapitulate disease pathology more accurately than existing disease models when differentiated into affected cell types. Thus, successful modeling of muscular diseases requires efficient differentiation of hiPSCs into skeletal muscles. hiPSCs transduced with doxycycline-inducible *MYOD1* (*MYOD1*-hiPSCs) have been widely used; however, they require time- and labor-consuming clonal selection procedures, and clonal variations must be overcome. Moreover, their functionality to exhibit muscular contraction has never been reported. Here, we demonstrated that bulk *MYOD1*- hiPSCs established with puromycin selection, but not with G418 selection, showed high differentiation efficiency, generating more than 80% Myogenin (MyoG)^+^ and Myosin heavy chain (MHC)^+^ muscle cells within seven days. Interestingly, bulk *MYOD1*-hiPSCs exhibited average differentiation properties compared with those of clonally established *MYOD1*- hiPSCs, suggesting that the bulk method may minimize the effects of clonal variations. Finally, three-dimensional muscle tissues were fabricated from bulk *MYOD1*-hiPSCs, which exhibited contractile force upon electrical pulse stimulation, indicating their functionality. Together, the findings indicate that our bulk differentiation requires less time and labor than existing methods, efficiently generates contractible skeletal muscles, and facilitates the generation of muscular disease models.

**Graphical Abstract:** 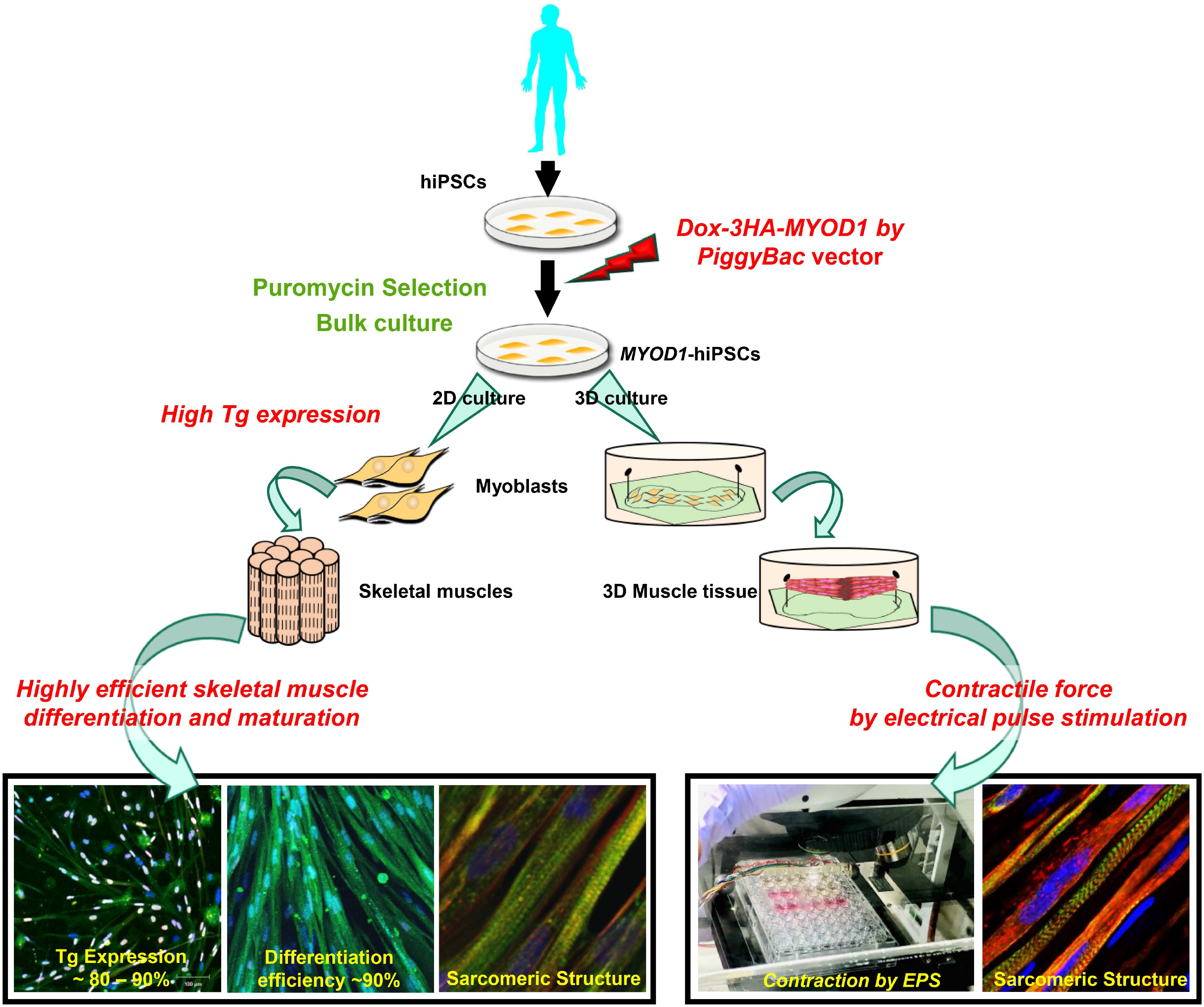

## Introduction

Human induced pluripotent stem cells (hiPSCs) are able to differentiate into any cell type in the body and have potential for use in clinical and research applications, such as cell-based therapies, human disease modeling, and drug discovery (1, 2). As disease-specific hiPSCs can differentiate into the affected cell types of patients, they can potentially recapitulate disease pathology more accurately than existing disease models, such as cell lines or mouse models, and are expected to overcome difficulties associated with species differences (3). However, successful modeling of human diseases with disease-specific hiPSCs requires efficient differentiation of hiPSCs into the affected cell types. Therefore, for efficient modeling of muscular diseases, it is crucial to develop a method for rapid, efficient, and reproducible differentiation of hiPSCs into skeletal muscles.

In the last several years, a variety of protocols for skeletal muscle differentiation from human pluripotent stem cells (hPSCs), including human embryonic stem cells (hESCs) and hiPSCs, have been reported (4). Compared with other strategies, recapitulation of skeletal muscle development *in vitro* with various morphogens and small molecules allows hPSCs to differentiate into skeletal muscles in a manner mimicking the process of *in vivo* development and induces the formation of skeletal muscles with more accurate physiological properties. Therefore, derived skeletal muscles may be applicable for mechanistic analysis of human skeletal muscle development and for regenerative therapies. However, in most cases, these differentiation protocols involve many complicated steps that take time and labor, and the efficiency of the differentiation is rather low (5–10). In contrast, forced expression of transcription factors associated with skeletal muscle development, such as *PAX3* and *PAX7*, which are expressed in satellite cells, or *MYOD1*, a master gene for muscular differentiation that induces muscular transdifferentiation of fibroblasts (11), more directly induces hPSCs to differentiate into skeletal muscles (12–18). Since these transgene-based methods may not allow physiological differentiation, skipping important developmental steps, skeletal muscles differentiated by these methods may have limited applications for analyses of skeletal muscle development and the pathogeneses of muscular diseases caused by developmental abnormalities and for regenerative therapies. However, skeletal muscle differentiation via the expression of transcription factors, especially *MYOD1*, is a valuable tool for the pathophysiological analysis of muscular diseases using disease-specific hiPSCs because it is highly efficient and rapid (19).

To date, success in skeletal muscle differentiation via *MYOD1* expression has been reported with the use of lentiviruses, adenoviruses, and *piggyBac* transposon vectors, and this method is widely used for analyses of muscular diseases (4, 14–18, 20, 21). As one of these tools, the *piggyBac* transposon vector for inducible expression of *MYOD1* is transduced into hPSCs, and the cells are selected with G418 or puromycin and screened for the appropriate *MYOD1*-hPSC clones exhibiting highly efficient skeletal muscle differentiation (16, 22). This differentiation method is relatively simple but still requires a time- and labor-consuming procedure to select appropriate *MYOD1*-hPSC clones in order to achieve highly efficient differentiation, and the proportion of appropriate *MYOD1*-hPSC clones is still relatively low. Clonal variations among *MYOD1*-hiPSC clones must also be considered (23), which may result in variation in the differentiation efficiency and the variability of the phenotypes, potentially masking pathologies or drug efficacies in analyses using disease-specific hPSCs. Therefore, multiple *MYOD1*-hPSC clones for each hPSC clone should be analyzed to confirm the reproducibility of the results. Notably, there are increasingly high expectations for analyses of sporadic diseases using disease-specific hiPSCs, such as analyses using samples from patient registries. In these cases, very large numbers of hiPSCs established from large numbers of patients and controls should be examined, but the establishment of multiple *MYOD1*-hiPSC clones for these cells is not realistic.

Based on these considerations, bulk establishment of *MYOD1*-hiPSCs or bulk differentiation of muscle cells from hiPSCs has been proposed to be beneficial for saving time and labor, minimizing the effects of clonal variations, and enabling quick analyses of relatively large numbers of hiPSCs. Through such a protocol, unknown pathological mechanisms may be clearly visualized. However, the bulk muscular differentiation reported thus far, achieved via expression of transcription factors using lentiviruses, adenoviruses, or *piggyBac* transposon vectors, requires complicated culture procedures. For example, it requires the induction of intermediate mesodermal progenitors and/or purification of transduced cells or muscular progenitors by flow cytometry and cell sorting (fluorescence activated cell sorting; FACS) using surface markers or fluorescent reporters to achieve high differentiation efficiency; otherwise, it achieves only moderate differentiation efficiency, inducing only 40% Myogenin (MyoG)^+^ cells and 60 to 70% Myosin heavy chain (MHC)^+^ cells (13-15, 17, 18, 20, 24, 25). Cell-to-cell variability in transgene expression may result in nonuniform differentiation, which may interfere with pathological analyses. Moreover, the functionality of hPSC-derived myotubes has been demonstrated primarily by patch clamp analysis or calcium imaging (13, 18, 19, 26, 27); rarely has it been demonstrated by the contraction of muscle tissues. In fact, the contractibility of hiPSC-derived muscle tissues differentiated via *MYOD1* expression has never been reported.

In this study, we demonstrated a simple and highly efficient method for contractible skeletal muscle differentiation from hiPSCs that takes advantage of bulk-established *MYOD1*-hiPSCs selected with puromycin. The relevance of bulk culture of *MYOD1*-hiPSCs was confirmed by comparing the muscular differentiation properties of the bulk-cultured cells with those of multiple clones of *MYOD1*-hiPSCs of the same origin. We also fabricated 3D muscle tissues from bulk *MYOD1*-hiPSCs, which exhibited contractile force upon electrical pulse stimulation. These results suggest that our bulk differentiation system facilitates the generation of disease models for analyses of the pathophysiological mechanisms of muscular disorders.

## Results

### Establishment of clonal or bulk *MyoD1*-hiPSCs with G418 or puromycin selection

To investigate whether we could efficiently derive skeletal muscles from bulk culture of *MYOD1*-hiPSCs without clonal selection and to determine how different selection markers, namely, G418 and puromycin, affect the muscular differentiation efficiency of *MYOD1*- hiPSCs, both bulk and clonal *MYOD1*-hiPSCs were established through transduction of a *piggyBac* transposon vector for the doxycycline (Dox)-inducible expression of human *MYOD1* tagged to *3 x HA* (*Dox-3HA-hMYOD1*) (with a selection cassette for G418 or puromycin) into hiPSCs (201B7) (2) (PB-TA-*3HA-hMYOD1*-ERN or PB-TA-*3HA- hMYOD1*-ERP2 (Fig. 1A, B). Ultimately, one bulk *MYOD1*-hiPSC line and six *MYOD1*- hiPSC clones were established each by G418 selection and puromycin selection (resulting in a G418-bulk line, a Puro-bulk line, G418-clones, and Puro-clones). All the established bulk and clonal *MYOD1*-hiPSCs were morphologically well maintained in the undifferentiated state and exhibited characteristic hESC-like morphology. We also confirmed that they expressed the hPSC markers Oct3/4 and Nanog by immunocytochemical (ICC) analysis, suggesting that all the bulk and clonal *MYOD1*-hiPSCs retained pluripotency (Fig. 1C and Supplementary Fig. S1).

**Fig. 1.**
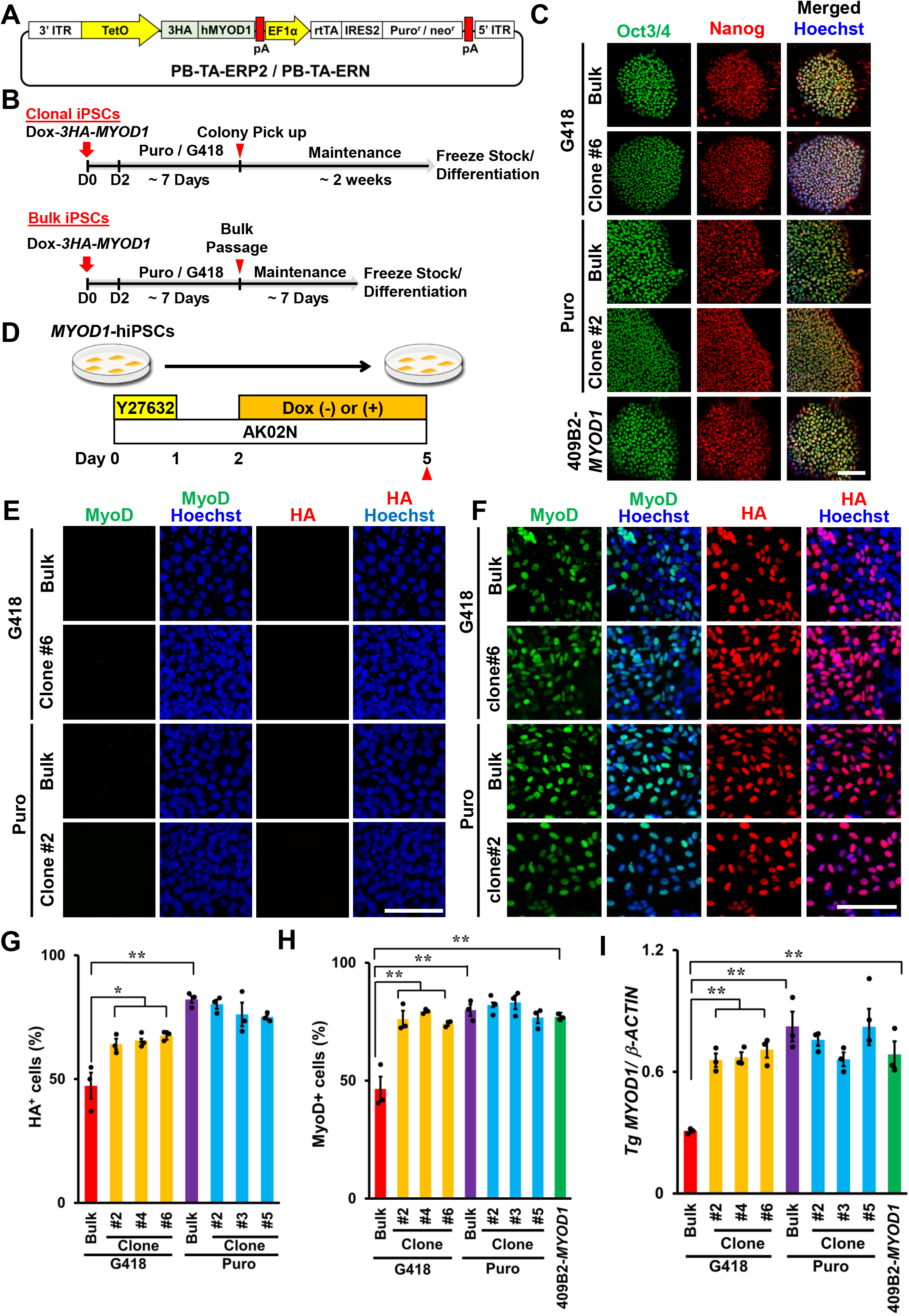
Establishment of bulk and clonal *MYOD1*-hiPSCs transduced with Dox-inducible *3HA-hMYOD1* (*Dox-3HA-MYOD1*). A. PiggyBac vector for Dox-inducible *3HA-hMYOD1* expression with the selection cassette for G418 or puromycin (*Dox-3HA-MYOD1*). B. Schematic of the establishment of bulk and clonal *MYOD1*-hiPSCs. The upper panel indicates the protocol for clonal *MYOD1*-hiPSCs, and the lower panel indicates the protocol for bulk *MYOD1*-hiPSCs. C. ICC analysis of the established *MYOD1*-hiPSCs for pluripotent stem cell markers (Oct3/4 and Nanog). The nuclei were stained with Hoechst 33258. All bulk and clonal *MYOD1*- hiPSCs retained the expression of pluripotent stem cell markers. Scale bar, 50 μm. D. Schematic of the analysis of transgene expression in the undifferentiated condition with or without Dox. *MYOD1*-hiPSCs were cultured in the presence of Dox from day 2 to day 5 or in the absence of Dox in the undifferentiated condition. Samples were collected at the times indicated by the closed triangles. E-H. ICC analysis of the expression of transgenes (HA) and MyoD1 in cells cultured without Dox (E) or with Dox (F). Scale bar, 200 μm. Quantitative analyses of HA^+^ and MyoD1^+^ cells are shown in G and H, respectively. The G418-bulk line exhibited lower expression of transgenes (HA) and MyoD1 than the G418-clones, whereas the Puro-bulk line showed expression of transgenes (HA) and MyoD1 similar to that of the Puro-clones. I. Transgene expression in bulk and clonal *MYOD1*-hiPSCs established with G418 or puromycin selection in the presence of Dox as examined by qRT-PCR and compared with that in control 409B2-*MYOD1*-hiPSCs. The amount of cDNA was normalized to that of human-specific *β-ACTIN*. The data are presented as the mean ± SEM, n = 3. *, *p < 0.05*, **, *p < 0.01*. ANOVA followed by *post hoc* Bonferroni test.

### Dox-inducible expression of *3HA-hMYOD1* transgenes in undifferentiated and differentiating conditions

To confirm Dox-inducible expression of the *3HA-hMYOD1* transgenes, 1.5 µg/ml Dox was added two days after the passage of all the *MYOD1*-hiPSC clones and bulk lines, and the cells were maintained in an undifferentiated condition for three days. Next, the expression of transgenes in the cells was analyzed in the presence and absence of Dox (Fig. 1D). ICC analysis revealed that in the presence of Dox, the proportions of HA^+^ and MyoD1^+^ cells in the G418-bulk line were 47.3 ± 5.3% and 46.5 ± 5.2%, respectively, whereas those in the G418-clones were significantly higher at approximately 65% (64.1 ± 2.2 to 67.6 ± 1.7%) and 75% (74.3 ± 1.1 to 79.5 ± 0.5%), respectively (Fig. 1F, G, H). In contrast, the proportions of HA^+^ and MyoD1^+^ cells in the Puro-bulk line were 82.2 ± 1.4% and 80.0 ± 2.5%, respectively, and were not significantly different from those in the Puro-clones, of which the proportions of HA^+^ and MyoD1^+^ cells were both approximately 80% (75.1 ± 1.0 to 80.3 ± 1.9% and 76.8 ± 2.4 to 83.2 ± 2.9%, respectively) (Fig. 1F, G, H). In the absence of Dox, the expression of the transgenes (HA) and MyoD1 could not be detected in any of the *MYOD1*-hiPSC clones or bulk lines (Fig. 1E). Dox-inducible expression of the transgenes was also examined by quantitative RT-PCR (qRT-PCR) in comparison with that in control 409B2-*MYOD1*-hiPSCs (409B2*-MYOD1*) that had been clonally established with PB-TAG- *hMYOD1*-ERP (puromycin selection, without the 3 x HA tag) (22) and were shown to differentiate into skeletal muscles with high efficiency. Consistent with the results of the ICC analysis, in the presence of Dox, the expression of transgenes in the G418-bulk line was significantly lower than that in the G418-clones and 409B2-*MYOD1*-hiPSCs (approximately 50% of the G418-clones and 409B2-*MYOD1*-hiPSCs) (Fig. 1I). In contrast, the Puro-bulk line expressed the same level of transgenes as Puro-clones and 409B2-*MYOD1*-hiPSCs, and the transgene expression levels in these *MYOD1*-hiPSCs in the presence of Dox were significantly higher than those in the G418-bulk line (Fig. 1I). Taken together, these results suggest that in undifferentiated states, the G418-bulk line was not capable of expressing sufficient transgenes in response to Dox, unlike the G418-clones and 409B2-*MYOD1*-hiPSCs. In contrast, Puro-bulk *MYOD1*-hiPSCs were able to express sufficient amounts of transgenes in response to Dox, comparable to the expression in the Puro-clones and 409B2-*MYOD1*- hiPSCs.

Considering the silencing of transgenes during differentiation, Dox-inducible transgene expression was also evaluated in differentiating hiPSCs (Fig. 2A). *MYOD1*-hiPSCs were differentiated into skeletal muscles as reported previously with modifications (16). From day 2 of differentiation, *MYOD1*-hiPSCs were cultured in the presence or absence of 1.5 μg/ml Dox for 3 days to induce the expression of *3HA-hMYOD1* (Fig. 2A). The transgene and endogenous *hMYOD1* expression was then analyzed by ICC analysis and qRT-PCR. Consistent with the results obtained in the undifferentiated condition, in response to Dox, the cells derived from the G418-bulk line were composed of only approximately 50% HA^+^ or MyoD1^+^ cells (53.9 ± 3.4% for HA, 48.0 ± 6.0% for MyoD1), whereas those derived from the G418-clones were composed of more than 75% HA^+^ or MyoD1^+^ cells (74.2 ± 2.5% to 80.2 ± 4.8% for HA, 75.3 ± 3.7% to 81.7 ± 4.9% for MyoD1), which was comparable to the results obtained with 409B2-*MYOD1*-hiPSCs for MyoD1 expression (77.5 ± 0.7%) (Fig. 2C- E). In contrast, the Puro-bulk line produced HA^+^ and MyoD1^+^ cells (80.6 ± 1.9% and 85.2 ± 2.3%, respectively) to the same extent as the Puro-clones, which yielded approximately 80% HA^+^ and MyoD1^+^ cells (71.4 ± 4.1% to 84.9 ± 3.0% for HA, 76.9 ± 0.6% to 87.3 ± 3.1% for MyoD1). This result was relatively similar to the MyoD1 result for the 409B2-*MYOD1*- hiPSCs (Fig. 2C-E). In the absence of Dox, the expression of transgenes (HA) and MyoD1 was not detected in any of the *MYOD1*-hiPSC bulk lines or clones (Fig. 2B). Similar results were obtained by qRT-PCR analysis. In response to Dox, the G418-bulk line expressed transgenes at only approximately 25% of the levels in G418-clones or 409B2-*MYOD1*- hiPSCs, in contrast to the Puro-bulk line, which expressed transgenes at a level comparable to those in the Puro-clones and 409B2-*MYOD1*-hiPSCs (Fig. 2F). Thus, in the differentiating condition, the G418-bulk line did not express *3HA-hMYOD1* transgenes sufficiently in response to Dox, whereas the Puro-bulk line, as well as the G418-clones and the Puro-clones, was capable of expressing higher levels of transgenes. Because sufficient expression of transgenes is required for efficient differentiation into skeletal muscles in this system, these results suggest that the selection of appropriate *MYOD1*-hiPSC clones is essential when *MYOD1*-hiPSCs are established with G418 selection, whereas clonal selection may not be required when the *MYOD1*-hiPSCs are established with puromycin selection.

**Fig. 2.**
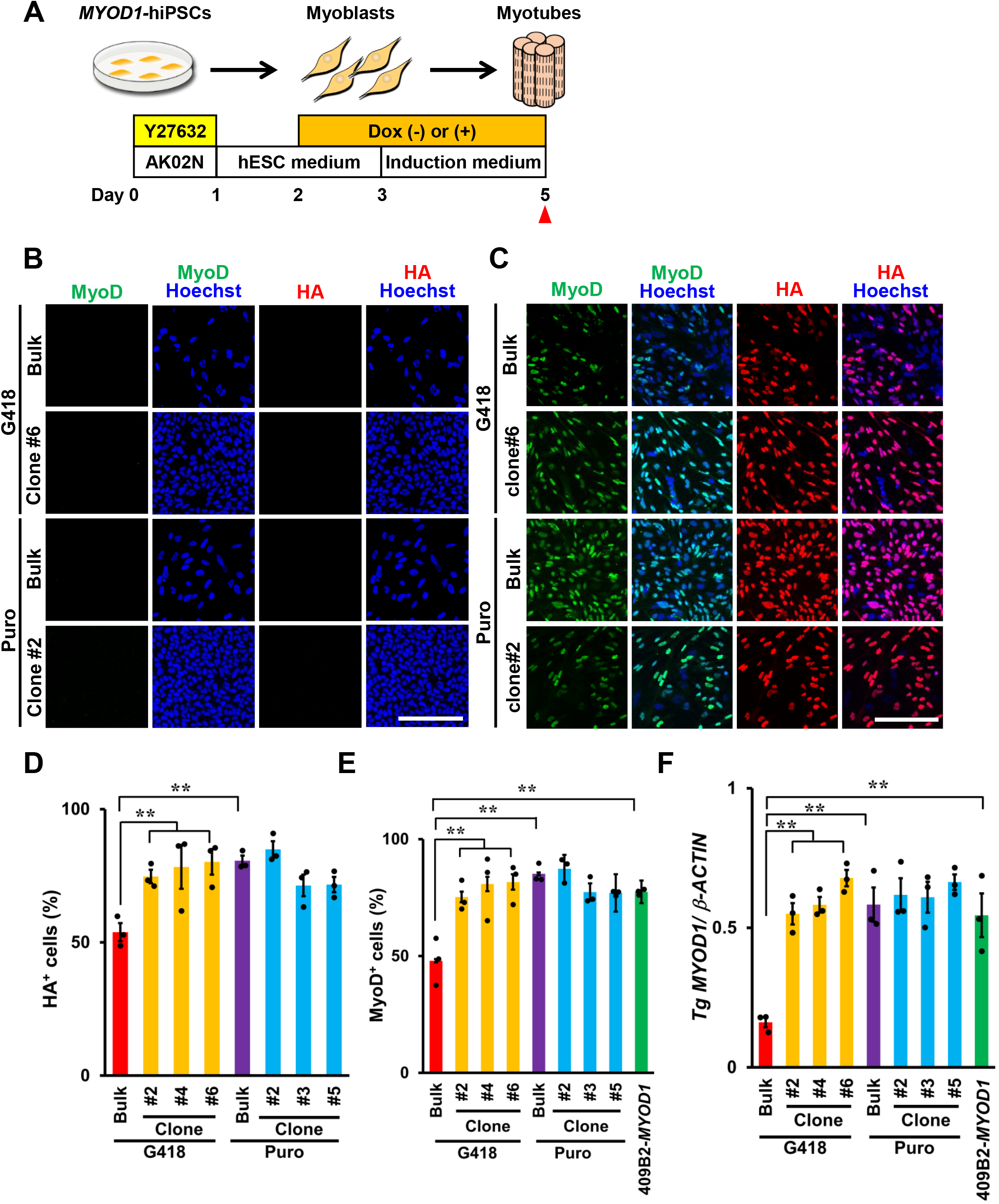
Dox-inducible expression of *3HA-hMYOD1* in the differentiating condition revealed similar transgene expression in Puro-bulk *MYOD1*-hiPSCs and clonal *MYOD1*- hiPSCs, but not G418-bulk *MYOD1*-hiPSCs. A. Schematic of the analysis of transgene expression in the differentiating condition with or without Dox. *MYOD1*-hiPSCs were induced to differentiate into skeletal muscles in the presence of Dox from day 2 to day 5 or cultured in the absence of Dox. Samples were collected at the times indicated by the closed triangles. B-E. ICC analysis of the expression of transgenes (HA) and MyoD1 in cells cultured in the differentiating condition without Dox (C) or with Dox (D). Scale bar, 200 μm. Quantitative analyses of HA^+^ and MyoD1^+^ cells are shown in D and E, respectively. The expression levels of transgenes (HA) and MyoD1 were lower in the G418-bulk line than in the G418-clones but were similar in the Puro-bulk line and the Puro-clones. F. Transgene expression in bulk and clonal *MYOD1*-hiPSCs established with G418 or puromycin selection in the differentiating conditions in the presence of Dox as examined by qRT-PCR and compared with that of control 409B2-*MYOD1*-hiPSCs. The amount of cDNA was normalized to that of human-specific *β-ACTIN*. The G418-bulk line expressed transgenes at only approximately 25% of the level observed in the G418 clones or 409B2-*MYOD1*- hiPSCs. The data are presented as the mean ± SEM, n = 3. *, *p < 0.05*, **, *p < 0.01*. ANOVA followed by *post hoc* Bonferroni test.

### Bulk *MYOD1*-hiPSCs established with puromycin selection were able to efficiently differentiate into mature skeletal muscles as clonally established *MYOD1*-hiPSCs

To confirm the correlation between transgene expression and the potential to differentiate into skeletal muscles, all bulk *MYOD1*-hiPSCs and clonal *MYOD1*-hiPSCs were differentiated into myotubes to evaluate differentiation potential and myotube maturation (Fig. 3A). As expected, two days after Dox withdrawal (day 7), ICC analysis revealed that the G418-bulk line, which failed to sufficiently express *3HA-hMYOD1* transgenes in response to Dox, showed inefficient differentiation into myotubes, as indicated by the significantly lower proportions of MyoG^+^ cells (71.2 ± 3.3%), MHC^+^ nuclei (58.6 ± 3.8%), and MHC^+^ area (64.2 ± 2.6%) than in the G418-clones (MyoG^+^ cells: 90.2 ± 1.7% to 91.6 ± 0.8%, MHC^+^ nuclei: 78.6 ± 5.5% to 88.2 ± 0.9%, MHC^+^ area: 79.5 ± 4.2% to 85.4 ± 1.5%) and control 409B2-*MYOD1*-hiPSCs (MyoG^+^ cells: 91.6 ± 0.8%, MHC^+^ nuclei: 78.6 ± 5.5%, MHC^+^ area: 80.0 ± 2.0%) (Fig 3B-E). On the other hand, the Puro-bulk line exhibited 91.6 ± 0.4% MyoG^+^ cells, 87.6 ± 1.1% MHC^+^ nuclei, and 83.8 ± 1.4% MHC^+^ area after differentiation, which were comparable to those of the Puro-clones (MyoG^+^ cells: 90.4 ± 0.2% to 91.8 ± 0.8%, MHC^+^ nuclei: 87.3 ± 0.3% to 88.4 ± 1.4%, MHC^+^ area: 82.0 ± 2.2% to 85.5 ± 2.4%) and control 409B2-*MYOD1*-hiPSCs. These results suggest that Puro-bulk *MYOD1*-hiPSCs, which expressed transgenes at sufficient levels, were able to differentiate into skeletal muscles as efficiently as clonally established *MYOD1*-hiPSCs, but G418-bulk *MYOD1*- hiPSCs were not able to do so because they lacked sufficient transgene expression. Moreover, quantitative analysis of the parameters of mature myotubes, including myotube thickness, the number of nuclei per myotube, and the MHC^+^ area per myotube, revealed that the G418-bulk line exhibited significantly poorer maturation according to all the parameters (myotube thickness: 11.7 ± 0.1 μm, MHC^+^ nuclei per myotube: 2.0 ± 0.02, MHC^+^ area per myotube: 2333.3 ± 137.2 μm^2^) than G418-clones or control 409B2-*MYOD1*-hiPSCs (myotube thickness: 13.4 ± 0.3 μm to 13.5 ± 0.1 μm, MHC^+^ nuclei per myotube: 2.6 ± 0.1 to 2.9 ± 0.1, MHC^+^ area per myotube: 3283.1 ± 77.3 μm^2^ to 3960.0 ± 75.8 μm^2^). In contrast, the Puro-bulk line gave rise to mature skeletal muscles (myotube thickness: 13.9 ± 0.4 μm, MHC^+^ nuclei per myotube: 3.0 ± 0.1, area per myotube: 3604.8 ± 35 μm^2^ MHC^+^) as efficiently as the Puro-clones (myotube thickness: 13.9 ± 0.2 μm to 13.4 ± 0.4 μm, MHC^+^ nuclei per myotube: 2.8 ± 0.5 to 3.0 ± 0.4, MHC^+^ area per myotube: 3440.7 ± 173.2 μm^2^ to 3742.5 ± 31.8 μm^2^) and 409B2-*MYOD1*-hiPSCs (Fig. 3B, F-H). These results suggest that puromycin selection for the establishment of *MYOD1*-hiPSCs facilitates higher transgene expression than G418 selection, which is critically correlated with the efficiency of muscular differentiation of *MYOD1*-hiPSCs as well as maturation of iPSC-derived myotubes.

**Fig. 3.**
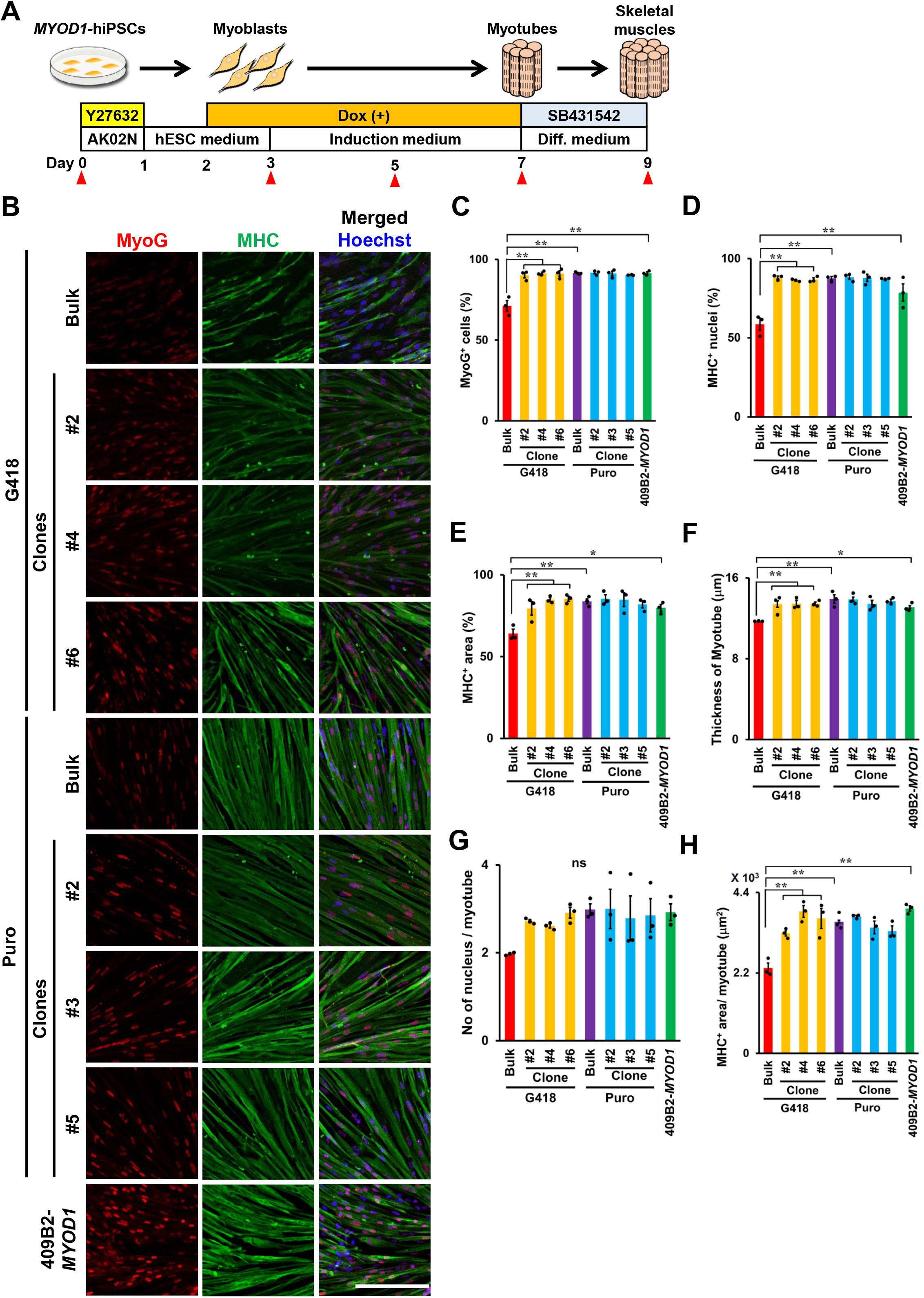
Bulk *MYOD1*-hiPSCs established with puromycin selection achieved efficient muscular differentiation and myotube maturation similar to that achieved by clonally established *MYOD1*-hiPSCs. A. Schematic presentation of skeletal muscle differentiation from *MYOD1*-hiPSCs. Samples were collected at the times indicated with the closed triangles. B. ICC analysis of myotubes derived from bulk and clonal *MYOD1*-hiPSCs established with G418 or puromycin selection and control 409B2-*MYOD1*-hiPSCs for the expression of MyoG and MHC at day 9 of differentiation. The nuclei were stained with Hoechst 33258. Scale bar, 200 μm. C-E. Quantitative analysis of the parameters for skeletal muscle differentiation, including the proportions of MyoG^+^ cells among total cells (C), the proportions of MHC^+^ nuclei among total nuclei (D), and the proportion of MHC^+^ area in the total area (E). Puro-bulk *MYOD1*- hiPSCs exhibited differentiation potential similar to that of clonal *MYOD1*-hiPSCs and control 409B2-*MYOD1*-hiPSCs, whereas G418-bulk *MYOD1*-hiPSCs exhibited lower differentiation potential than the other *MYOD1*-hiPSCs. F-H. Quantitative analysis of the parameters for myotube maturation, including the myotube thickness (F), number of nuclei per myotube (G), and MHC^+^ area per myotube (H). Puro- bulk *MYOD1*-hiPSCs exhibited myotube maturation potential similar to those of clonal *MYOD1*-hiPSCs and control 409B2-*MYOD1*-hiPSCs, whereas G418-bulk *MYOD1*-hiPSCs exhibited lower myotube maturation potential than the other *MYOD1*-hiPSCs. The data are presented as the mean ± SEM, n = 3. *, *p < 0.05*, **, *p < 0.01*. ANOVA followed by *post hoc* Bonferroni test.

### Bulk *MYOD1*-hiPSCs exhibited an average differentiation process compared to that of clonal *MYOD1*-hiPSCs and may have minimized the effects of clonal variations

To examine the differences between bulk and clonal *MYOD1*-hiPSCs and G418 and Puro selection in more detail, we further performed time-course analysis of skeletal muscle differentiation with all *MYOD1*-hiPSC bulk lines and clones. As differentiation progressed, all the bulk and clonal *MYOD1*-hiPSCs gradually showed morphological differentiation into myoblasts and subsequently into myotubes, and most of the cells formed myotube-like aligned structures by day 7, the time of Dox withdrawal. As expected, clonal *MYOD1*-hiPSCs and Puro-bulk *MYOD1*-hiPSCs exhibited further maturation of myotubes, whereas G418- bulk *MYOD1*-hiPSCs showed increased spaces among myotubes and decreased numbers of myotubes and gave rise to nonmuscle cells with a more flattened morphology (Fig. 4A, Supplementary Figs. S3, S4). Time-course gene expression analysis of markers associated with skeletal muscle development was also performed in the presence of Dox by qRT-PCR (Fig. 4B and Supplementary Fig. S5). The expression of the pluripotent stem cell markers, including *NANOG* and *OCT3/4*, gradually and similarly decreased in all *MYOD1*-hiPSCs over the course of differentiation regardless of the method used for *MYOD1*-hiPSC establishment. In all the bulk and clonal *MYOD1*-hiPSCs, the expression of the premyogenic marker CD56 and the core myogenic regulatory factor (MRF) total *MYOD1*, including endogenous *MYOD1* and transgenes, appeared on day 3, just after the induction of the transgenes and gradually increased to show the highest expression on day 5, after which it decreased. Endogenous *MYOD1* expression peaked at day 5, just after transgene induction, and was downregulated thereafter; however, its expression was maintained at approximately 50% of the peak expression even after the withdrawal of Dox at day 7. Other MRFs, including *MYOG* and *MYF6*, which are crucial for the fate determination and terminal differentiation of skeletal muscles (28, 29), showed gradual increases in their expression along with differentiation. *MYOG* showed decreased expression after the withdrawal of Dox at day 7, whereas *MYF6* showed continuous upregulation thereafter. The expression of *MYF5* was not detected at all, consistent with previous reports (14–16), probably due to negative feedback regulation from the overexpression of *MYOD1,* which has similar roles in muscular differentiation. With regard to the maturation of myotubes, a transcription factor critically involved in the maintenance of skeletal muscles, *MEF2C*; the mature skeletal muscle markers *MYH2* and *MYH7*; and the fusion marker *TMEM8C* showed gradual increases, with peak expression from day 5 to day 7, followed by decreases in their expression thereafter. The expression of mature skeletal muscle markers, except for *TMEM8C*, was maintained at approximately 50% of the peak expression even after the withdrawal of Dox at day 7, similar to pattern for endogenous *MYOD1*, suggesting that the expression of these molecules depends on *MYOD1* expression and may contribute to the continuous maturation of the cells at later stages of differentiation.

**Fig. 4.**
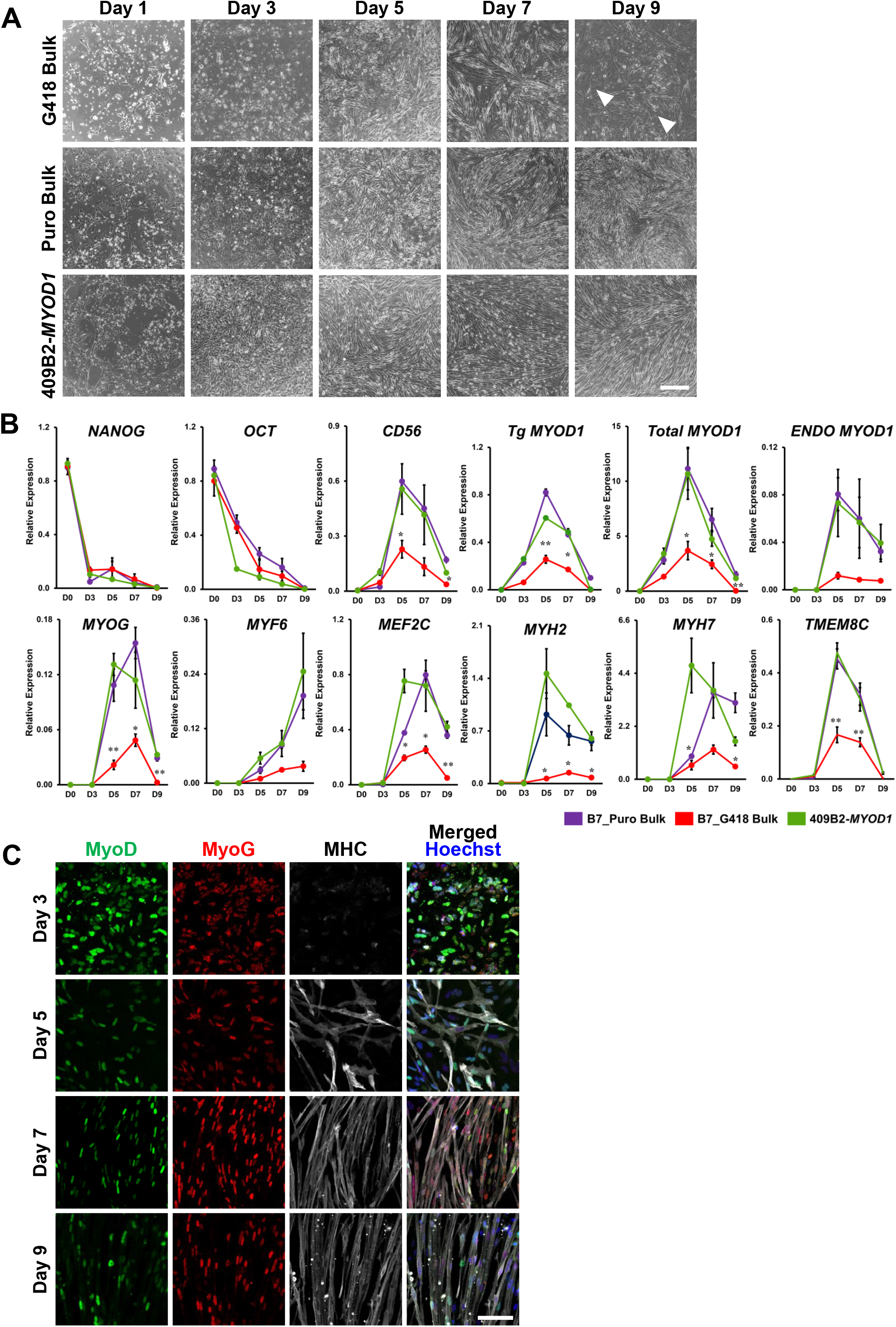
Time course analysis revealed highly efficient muscular differentiation of Puro- bulk *MYOD1*-hiPSCs. A. Brightfield images showing the time course of muscular differentiation of G418-bulk and Puro-bulk *MYOD1*-hiPSCs as well as control 409B2-*MYOD1*-hiPSCs. After day 7, G418- bulk *MYOD1*-hiPSCs featured non-muscle cells with flattened morphology and showed fewer myotubes and larger spaces among myotubes than did Puro-bulk and clonal *MYOD1*-hiPSCs. Scale bar, 100 μm. B. Time course gene expression analysis of G418-bulk, Puro-bulk, and control 409B2- *MYOD1*-hiPSCs along with muscular differentiation. The amount of cDNA was normalized to that of human-specific *β-ACTIN* and is presented as the relative expression in undifferentiated hiPSCs (*NANOG* and *OCT3/4*), the human myoblast cell line Hu5/KD3 differentiated for 3 days (*CD56*, total and endogenous *MYOD1*, *MYOG*, *MYF6*, *MEF2C*, *MYH2*, *MYH7*, and *TMEM8C*), and EKN3-*MYOD1*-hiPSCs differentiated for 5 days (*Tg MYOD1*). The data are presented as the mean ± SEM, n = 3. *, *p < 0.05*, **, *p < 0.01 vs.* 409B2-*MYOD1*-hiPSCs. ANOVA followed by *post hoc* Bonferroni test. C. Time course ICC analysis of Puro-bulk *MYOD1*-hiPSCs for MyoD, MyoG, and MHC along with muscular differentiation. The number of MyoD^+^ cells gradually decreased after day 7, while MHC^+^ cells appeared around day 5 and formed aligned myotubes from day 7. Scale bar, 100 μm.

When bulk and clonal *MYOD1*-hiPSCs and G418 and puromycin selection were compared, the Puro-bulk line showed expression similar to that of the Puro-clones and 409B2-*MYOD1*-hiPSCs, whereas the G418-bulk line showed lower expression levels than other *MYOD1*-hiPSCs throughout the differentiation process. These findings are consistent with the data showing the poor differentiation potential of the G418-bulk line. More importantly, in contrast to the large clonal variations observed in differentiating Puro-clones, the Puro-bulk line showed average expression of genes associated with skeletal muscle differentiation among the six Puro-clones or similar expression to that in six G418-clones during the differentiation processes (Supplementary Fig. S5). Considering the average transgene expression in undifferentiated and differentiating conditions and the average values of the parameters of skeletal muscle differentiation and maturation in the Puro-bulk line among Puro-clones and G418-clones (Fig. 1F-I, Fig. 2C-F, Fig. 3B-H), these results suggest that Puro-bulk *MYOD1*-hiPSCs have average properties of clonal *MYOD1*-hiPSCs. This averaging may minimize the effects of clonal variation among *MYOD1*-hiPSC clones, which is responsible for a bottleneck in the analysis of disease-specific hiPSCs.

Finally, we performed time-course ICC analysis of the differentiation of Puro-bulk *MYOD1*-hiPSCs to determine the differentiation potential of the cells. We found that the expression of MyoG was first observed at day 3 and was followed by the expression of MHC at day 5. Morphological alteration into aligned myotubes was observed from day 7 of differentiation, which was consistent with the results of time-course morphological and qRT-PCR analyses (Fig. 4A-C). Taken together, these results suggested that bulk *MYOD1*-hiPSCs established by puromycin selection were capable of differentiating into mature skeletal muscles as efficiently as clonally established *MYOD1*-hiPSCs.

### Reproducibility of efficient muscular differentiation in various hiPSC clones using Puro-bulk *MYOD1*-hiPSCs

To confirm the reproducibility of the differentiation potential of bulk cultures of *MYOD1*- hiPSCs generated via puromycin selection, three hiPSC clones (EKN-3, YFE-19, and TIGE-9) established from fibroblasts of adult healthy individuals (30–32) were transduced with the *Dox-3HA-hMYOD1 PiggyBac* vector with a puromycin selection cassette in combination with transposase and differentiated into skeletal muscles in bulk following the same protocol (Fig. 1B and 3A). After the transduction of *3HA-hMYOD1* and selection with puromycin, all three hiPSC clones maintained their pluripotent state (Fig. 5A). ICC analysis at day 9 of differentiation revealed that all of the Puro-bulk *MYOD1*-hiPSCs efficiently differentiated into skeletal muscles, as confirmed by the proportions of MyoG^+^ cells, MHC^+^ nuclei, and MHC^+^ areas of more than 90%, 85%, and 80%, respectively (Fig. 5B-E). Quantitative analysis of the myotube thickness, numbers of nuclei per myotube, and MHC^+^ area per myotube revealed that all the Puro-bulk *MYOD1*-hiPSCs exhibited a potential to differentiate into mature myotubes (myotube thickness: 15.0 ± 0.3 μm to 16.1 ± 0.4 μm, MHC^+^ nuclei per myotube: 2.1 ± 0.1 to 2.8 ± 0.2, MHC^+^ area per myotube: 3177.7 ± 64.9 μm^2^ to 3532.5 ± 176.1 μm^2^) similar to that of 201B7-*MYOD1*- and 409B2-*MYOD1*-hiPSCs (Fig. 3B-H, and Fig. 5B-D). Moreover, time course analysis of the expression of myogenic markers by qRT-PCR analysis revealed that all the Puro-bulk *MYOD1*-hiPSCs exhibited gene expression profiles similar to those observed in Puro-bulk 201B7*-MYOD1-* and control 409B2-*MYOD1*-hiPSCs (Figs. 4B, 5E, and Supplementary Fig. S5). Finally, by ICC analysis of α-Actinin and Titin at day 9 of differentiation, sarcomere formation was clearly visualized in myotubes derived from all Puro-bulk *MYOD1*-hiPSCs, suggesting that these hiPSC-derived myotubes were mature enough to exhibit functional properties (Fig. 5F) (33). These results suggest that the highly efficient differentiation of Puro-bulk *MYOD1*-hiPSCs is reproducible and applicable to various hiPSCs for simple and efficient mature skeletal muscle differentiation.

**Fig. 5.**
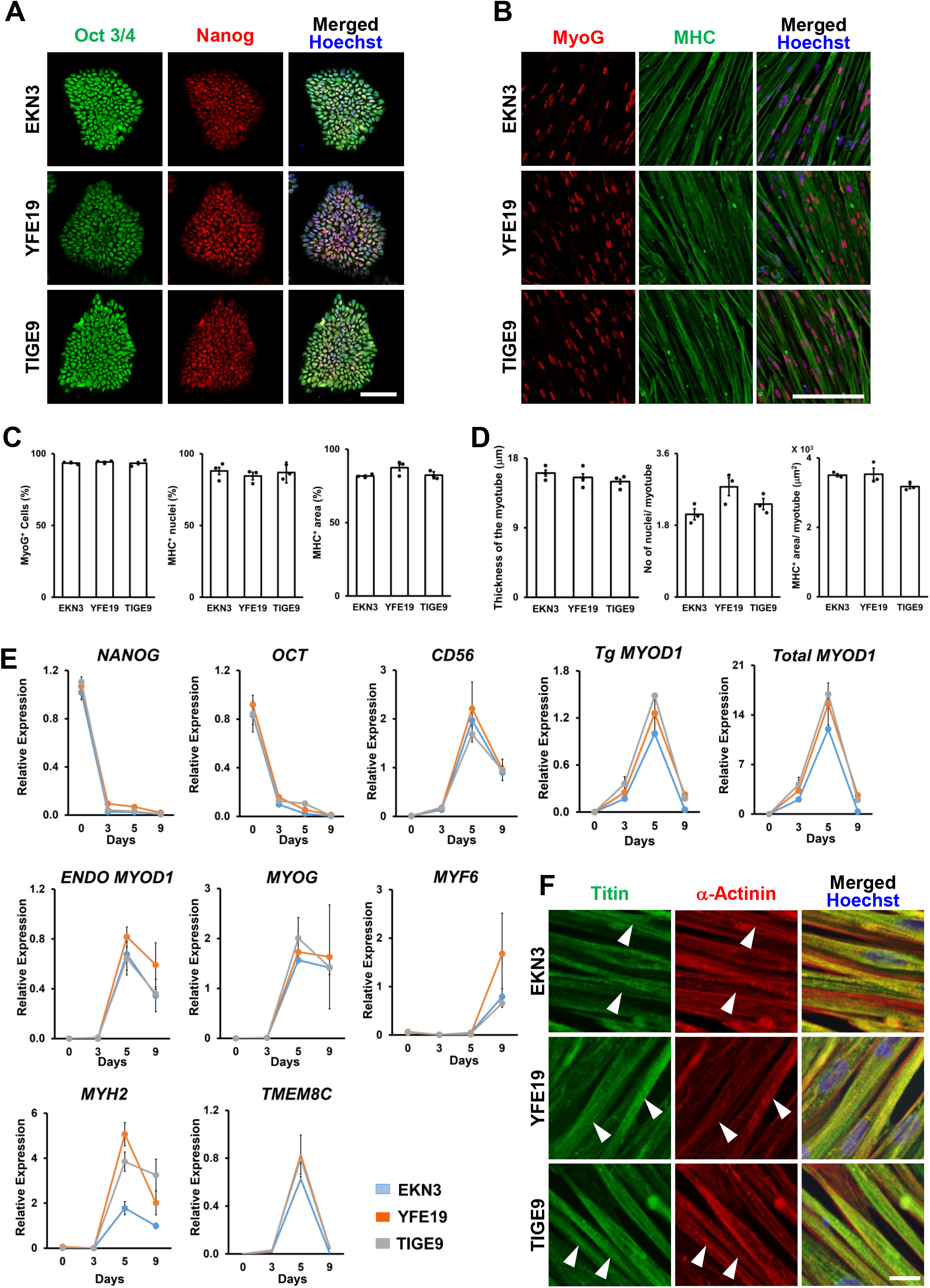
Reproducibility of highly efficient skeletal muscle differentiation in various hiPSCs. A. Establishment of *3HA-hMYOD1*-hiPSCs from three control iPSC clones, EKN3, YFE19, and TIGE9, by the Puro-bulk method. The pluripotency of established *MYOD1*-hiPSCs was confirmed by ICC analysis of Oct3/4 and Nanog. The nuclei were stained with Hoechst 33258. Scale bar, 100 μm. B. All three Puro-bulk *MYOD1*-hiPSC lines efficiently differentiated into skeletal muscles in the presence of Dox, as shown by ICC analysis of MyoG and MHC at day 9. C. Quantitative analysis of the parameters for skeletal muscle differentiation, including the proportions of MyoG^+^ cells among total cells, the proportions of MHC^+^ nuclei among total nuclei, and the proportion of MHC^+^ area in the total area. The data are presented as the mean ± SEM, n = 3. Scale bar, 200 μm. D. Quantitative analysis of the parameters for myotube maturation, including the myotube thickness, the number of nuclei per myotube, and the MHC^+^ area per myotube. The data are presented as the mean ± SEM, n = 3. E. Time course gene expression analysis of three Puro-bulk *MYOD1*-hiPSC lines established from EKN3, YFE19, and TIGE9 along with muscular differentiation. The amount of cDNA was normalized to that of human-specific *β-ACTIN* and is presented as the relative expression in undifferentiated hiPSCs (*NANOG* and *OCT3/4*), the human myoblast cell line Hu5/KD3 differentiated for 3 days (*CD56*, total and endogenous *MYOD1*, *MYOG*, *MYF6*, *MEF2C*, *MYH2*, and *TMEM8C*), and EKN3-*MYOD1* iPSCs differentiated for 5 days (*Tg MYOD1*). The data are presented as the mean ± SEM, n = 3. F. ICC analysis indicating the maturation of myotubes according to the sarcomere structure as shown by Titin and α-Actinin staining. The nuclei were stained with Hoechst 33258. The arrowheads indicate sarcomere formation. Scale bar, 20 μm.

### 3D muscle tissues were fabricated from Puro-bulk *MYOD1*-hiPSCs and exhibited contractility upon electrical pulse stimulation

To investigate the functionality of skeletal muscles derived from bulk *MYOD1*-hiPSCs, 3D muscle tissues were fabricated from Puro-bulk 201B7-*MYOD1*-hiPSCs and control 409B2-*MYOD1*-hiPSCs on microdevices and examined for their contractility (Fig. 6A). Both of these *MYOD1*-hiPSCs were differentiated into skeletal muscle cells by the same monolayer differentiation protocol until day 3 and were then redissociated and processed for the fabrication of 3D muscle tissues in the microdevices. Both types of *MYOD1*-hiPSCs gave rise to similar 3D muscle tissues that expressed α-Actinin and Titin at day 17 of differentiation according to immunohistochemical (IHC) analysis, which uncovered sarcomere structures and uniform alignment of striated muscle fibers (Fig. 6B-E). On days 11, 13, 15, and 17, the contractile force of the fabricated muscle tissues was measured upon electrical pulse stimulation (EPS, 2 msec, 20 V, 30 Hz). On day 13 (day 10 of 3D culture), Puro-bulk 201B7-*MYOD1*-hiPSC- and 409B2-*MYOD1*-hiPSC-derived muscle tissues exhibited contractile forces of 1.7 ± 0.6 µN and 1.2 ± 0.1 µN, respectively, in response to EPS (Fig. 6F, G). On day 15 (day 12 of 3D culture), the contractile force of 201B7-*MYOD1*- hiPSC- and 409B2-*MYOD1*-hiPSC-derived muscle tissues increased to 4.3 ± 1.1 µN and 2.6 ± 0.5 µN, respectively, after which it decreased (Fig. 6F, G, H). To confirm the relevance of our 3D muscle cultures, 3D muscle tissues were also fabricated from mouse (C2C12) and human (Hu5/KD3) myoblast cell lines. The resulting tissues showed contractile forces of 6.5 ± 0.3 µN and 14.8 ± 2.0 µN, respectively, at day 6 of 3D culture, consistent with a previous report (34), and maximum contractile forces of 10.4 ± 1.7 µN and 23.4 ± 2.3 µN, respectively, at day 8 of 3D culture (Fig. 6H). Although the observed maximum contractile forces of iPSC-derived muscle tissues were still lower than those observed in C2C12 and Hu5/KD3-derived muscle tissues (Fig. 6H), they were comparable to those previously reported in hiPSC-derived 3D-muscle tissues fabricated using similar microdevices, which showed a maximum contractile force of 2.0 µN (35). More importantly, because the previous study showing the fabrication of 3D muscle tissues by *MYOD1* expression required a more complex protocol and did not demonstrate contractile force (36), our findings demonstrate, for the first time, that hiPSC-derived muscle tissues generated via expression of *MYOD1*, especially those generated by a simple method using Puro-bulk *MYOD1*-hiPSCs, can give rise to functional 3D muscle tissues with contractile force. This strategy may be a valuable tool for future analyses of muscular disease.

**Fig. 6.**
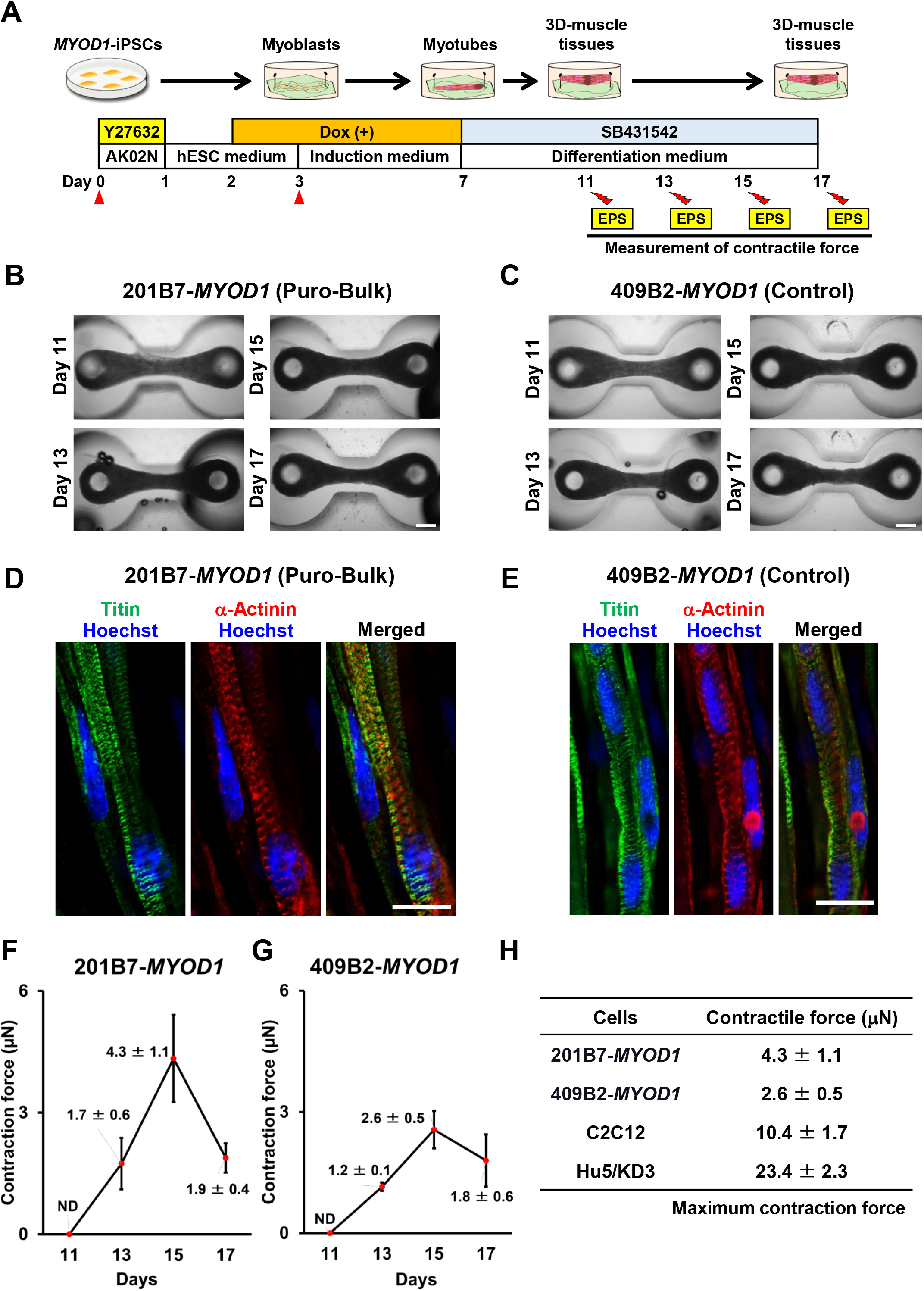
3D muscle tissues fabricated from Puro-bulk *MYOD1*-hiPSCs exhibited contractile force, indicating the functionality of the hiPSC-derived muscle tissues. A. Schematic of the fabrication of 3D muscle tissues. On day 3 of differentiation, differentiating cells were dissociated and replated on microdevices. On day 11, muscle tissues were pulled up at the top of the pillar and processed for measurement of contractile force elicited by electrical pulse stimulation (EPS) at days 11, 13, 15 and 17. B, C. Brightfield top-view images of muscle tissues derived from Puro-bulk 201B7-*MYOD1*- hiPSCs (B) and control 409B2-*MYOD1*-hiPSCs (C) at days 11, 13, 15, and 17. Scale bar, 500 μm. D, E. IHC analysis of fabricated muscle tissues from Puro-bulk 201B7-*MYOD1*-hiPSCs (D) and control 409B2-*MYOD1*-hiPSCs (E) for Titin and α-Actinin at day 17 of differentiation. Sarcomere formation was clearly observed in both muscle tissues. The nuclei were stained with Hoechst 33258. Scale bar, 20 µm. F, G. Measurement of the contractile force of the muscle tissues derived from Puro-bulk 201B7-*MYOD1*-hiPSCs (F) and control 409B2-*MYOD1*-hiPSCs (G) at days 11, 13, 15, and 17 of differentiation. The maximum contractile force is indicated at day 15. The data are presented as the mean ± SEM, n = 4. H. Comparison of the maximum contractile force of muscle tissues derived from Puro-bulk 201B7-*MYOD1*-hiPSCs, control 409B2-*MYOD1*-hiPSCs, C2C12, and Hu5/KD3 cells.

## Discussion

Our data suggest that for highly efficient skeletal muscle differentiation using bulk *MYOD1*-hiPSCs, the selection marker plays important roles in increasing transgene expression efficiency, which may ultimately result in increased differentiation efficiency. Although different selection markers are commonly speculated to affect transgene expression and the properties of transduced cells, there have been few reports describing the effects of selection markers, especially their effects on the differentiation properties of hPSCs. In our bulk culture differentiation, bulk *MYOD1*-hiPSCs established with puromycin selection showed similar or higher transgene expression and differentiation efficiency than clonally established *MYOD1*-hiPSCs, whereas G418-bulk *MYOD1*-hiPSCs exhibited poor transgene expression and differentiation potential (Fig. 1F-H, Fig. 2C-F, Fig. 3). Notably, if we picked appropriate clones, clonally established *MYOD1*-hiPSCs selected with either G418 or puromycin achieved almost the same transgene expression and differentiation potential as control 409B2-*MYOD1*-hiPSCs. These data suggest that puromycin selection has the advantage of efficient transgene expression even for bulk *MYOD1*-hiPSCs and that sufficient transgene expression is key for high differentiation efficiency. Interestingly, bulk *MYOD1*- hiPSCs established with a higher concentration of puromycin (1.5 μg/ml), in which higher transgene expression could be achieved, did not show significantly greater differentiation efficiency than those established with a lower concentration of puromycin (0.5 μg/ml) (data not shown). This finding suggests that not only the transgene expression level but also other factors, such as cell-to-cell variability in transgene expression, may affect the efficiency of muscular differentiation from bulk *MYOD1*-hiPSCs.

What is responsible for the differences between G418 and puromycin? Both G418 and puromycin are aminoglycoside antibiotics that are commonly used for the selection of mammalian cells transduced with transgenes. Both G418 and puromycin inhibit protein synthesis by suppressing translation at ribosomes and induce cell death. However, puromycin is distinct from G418 in that it induces cell death more quickly and powerfully (37, 38). In most cells, G418 exhibits relatively slow action, taking more than one week to gradually induce cell death, whereas puromycin may exhibit faster action to induce complete cell death within 2 to 4 days if the cells do not express resistance genes (37–39).

Moreover, a previous analysis using HEK293 cells showed distinct effects on the levels of and cell-to-cell variability in transgene expression (40). In that study, G418 was shown to induce lower expression of transgenes with higher cell-to-cell variability, whereas puromycin was shown to induce higher transgene expression (10 times more than G418) with lower cell-to-cell variability (40). This difference might have been caused by the different stabilities and activities of the resistance genes. If the selection cassette for G418, *neo^r^* from transposons (Tn601 or Tn5) that express aminoglycoside-3’-phosphotransferase (APH), is relatively stable and has relatively high antibiotic-inactivating activity, even the cells expressing relatively low levels of transgene and *neo^r^* may survive after G418 selection, and cells expressing the transgenes at a variety of levels may remain. On the other hand, if the selection cassette for puromycin, *pac* (encoding puromycin N-acetyl-transferase), is relatively unstable and has relatively low antibiotic-inactivating activity, only cells expressing relatively high levels of the transgene and *pac* may survive after puromycin selection, and the transgene expression levels of the remaining cells may be relatively high with reduced cell-to-cell variability. In the abovementioned analysis using HEK293 cells, the level and uniformity of transgene expression were shown to be the highest with zeocin, relatively high with puromycin and hygromycin, and the lowest with G418 and blasticidin S (40). These results suggest that for efficient transduction of genes into hPSCs, zeocin, puromycin, or hygromycin may be the best choices to obtain bulk cells with increased transgene expression with reduced cell-to-cell variability or to increase the possibility of picking clones with higher transgene expression.

In addition, we achieved quick and efficient differentiation of bulk *MYOD1*-hiPSCs into skeletal muscles, which may have minimized the effects of clonal variations. To date, several methods have been reported that achieve bulk differentiation of hiPSCs into skeletal muscles using lentiviruses, adenoviruses, or *piggyBac* transposon vectors (4, 14, 15, 18, 20, 25). However, most of these differentiation protocols have never achieved high differentiation efficiency, short culture periods, and simple culture methods simultaneously. Moreover, with these methods, researchers must start from undifferentiated and unmodified hiPSCs for every differentiation, which causes considerable differentiation variability. Importantly, previous studies have not compared bulk differentiation with clonal differentiation. In this study, bulk *MYOD1*-hiPSCs were established by a simple method: a single transfection of a *piggyBac* vector with transposase followed by selection with puromycin for seven days (Fig. 1B, lower protocol). The reproducibility of the differentiation from bulk *MYOD1*-hiPSCs was confirmed with at least four clones of hiPSCs established independently, and all the bulk *MYOD1*-hiPSCs achieved highly efficient skeletal muscle differentiation, resulting in more than 80% MyoG^+^ or MHC^+^ cells (Figs. 3-5). This percentage is significantly higher than those obtained with the previously reported bulk culture methods using *piggyBac* transposon vectors or adenoviruses, which have yielded approximately 40 to 50% MyoG^+^ cells and 50 to 70% MHC^+^ cells (14, 18, 25). Moreover, in our method, once we generated *MYOD1*-hiPSCs, we did not have to return to using unmodified, undifferentiated hiPSCs, suggesting that we can easily reproduce differentiation at any time. More importantly, we confirmed that the Puro-bulk *MYOD1*-hiPSCs showed gene expression and differentiation efficiency similar to those of previously confirmed control 409B2-*MYOD1*-hiPSCs and conventional clonal *MYOD1*-hiPSCs (Fig. 3, Supplementary Figs. S2-5). Interestingly, we found, for the first time, that Puro-bulk *MYOD1*-hiPSCs exhibited average properties of clonally established *MYOD1*-hiPSCs of the same origin, which showed considerable variability among clones. As our bulk differentiation method requires only one bulk *MYOD1*-hiPSC line for one hiPSC clone, it may save valuable time and labor, enable the analysis of large numbers of hiPSC clones within short periods, and provide valuable *in vitro* skeletal muscle models, especially for analyses using disease-specific hiPSCs. The advantage of bulk culture has also been shown in the establishment of disease-specific iPSCs of sporadic amyotrophic lateral sclerosis (ALS) and in an analysis of motor neurons derived from them. A collection of a large number of patient-derived lymphoblastoid cells from a multicenter ALS cohort in Japan, named the Japanese Consortium for Amyotrophic Lateral Sclerosis Research (JaCALS), was established for hiPSCs in a bulk manner. The cells were subsequently differentiated into motor neurons and exhibited differentiation efficiencies and phenotypes similar to those of corresponding clonally established hiPSCs. The phenotypes also reflected those observed in the corresponding ALS patients (41). Therefore, muscular differentiation of bulk *MYOD1*- hiPSCs with our system may be particularly beneficial for analyses of sporadic diseases, which require analysis of large numbers of patient-derived iPSCs.

Furthermore, we could have derived mature and functional skeletal muscles from bulk *MYOD1*-hiPSCs. Sarcomere formation was clearly shown by the expression of α-Actinin and Titin in monolayer myotubes, and 3D muscle tissues were fabricated from Puro-bulk *MYOD1*-iPSCs (8, 27, 33) (Fig. 5J, 6D, E). Moreover, the fabricated 3D-muscle tissue exhibited contractile force upon EPS (maximum contractile force of 4.3 μN; Fig. 6F-G) that was comparable to the contractile force observed in previously reported hiPSC-derived 3D- muscle tissues fabricated using similar microdevices (2.0 μN) (35), suggesting the functionality of the muscle tissues derived from bulk *MYOD1*-iPSCs. Notably, there have been only a few studies showing the fabrication of contractile 3D muscle tissues from hiPSCs (24, 35), and only one study using *MYOD1* expression; however, that study required a more complex protocol and did not show muscular contraction (36). Thus, this is the first report demonstrating the fabrication of contractile 3D muscle tissues via the expression of *MYOD1*, especially with a simplified method using bulk culture of *MYOD1*-hiPSCs. The observed contractile force was still lower than that observed in C2C12 or Hu5/KD3 cells (Fig. 6H) (34). In addition, the differentiation system still needs to be optimized to produce more mature and contractible muscle tissues for further analysis. Nevertheless, our skeletal muscle differentiation system using Puro-bulk *MYOD1*-hiPSCs will provide a new approach for disease-specific hiPSC studies and may facilitate the generation of better disease models for the pathophysiological analysis of muscular disorders.

## Materials and Methods

### Construction of a Dox-inducible *3HA-hMYOD1*-expressing *PiggyBac* vector

Human *MyoD1* (*hMYOD1*) cDNA was cloned and inserted into the pENTR vector and tagged with 3 x HA at the N-terminus. Then, *3HA-hMYOD1* cDNA was transferred to PB- TA-ERN (AddGene #80474) or PB-TA-ERP2 (AddGene #80477) (42) (kindly provided by Dr. Knut Woltjen, Kyoto University) with Gateway cloning technology to generate *piggyBac* vectors for Dox-inducible expression of *3HA-hMYOD1 (Dox-3HA-hMYOD1* vector), PB-TA- *3HA-hMYOD1*-ERN (for selection with G418) and PB-TA-*3HA-hMYOD1*-ERP2 (for selection with puromycin).

### Human iPSC culture and *MYOD1*-iPSC establishment

The hiPSCs used in this study were established previously (EKN3, YFE-19 and TIGE-9) (30–32) or kindly provided by Dr. Shinya Yamanaka, Kyoto University, Japan (201B7 and 409B2) (2, 43). The hiPSCs were grown on mitomycin C-treated SNL murine fibroblast feeder cells on gelatin-coated (0.1%) tissue culture dishes and were maintained in standard hESC medium consisting of Dulbecco’s modified Eagle’s medium (DMEM)/F12 supplemented with 20% KnockOut™ serum replacement (KSR; Thermo Scientific, USA), nonessential amino acids, 0.1 mM 2-mercaptoethanol (2-ME; Sigma-Aldrich, USA), and 4 ng/ml FGF-2 (PeproTech, USA) at 37 °C in a humidified atmosphere containing 3% CO2. For feeder-free culture of hiPSCs, hiPSCs maintained on SNL feeder cells were transferred to plates coated with iMatrix-511 silk (Nippi, Japan) and maintained in StemFit medium (AK02N, Ajinomoto, Japan) at 37 °C in a humidified atmosphere containing 5% CO2.

To generate *MYOD1*-hiPSCs, semiconfluent hiPSCs maintained under feeder-free conditions were dissociated into single cells by Accutase (12679-54, Nacalai Tesque, Japan), and 1.25 x 10^5^ cells were electroporated with 0.5 μg of PB-TA-*3HA-hMYOD1*-ERN or PB- TA-*3HA-hMYOD1*-ERP2 in combination with 0.5 μg of transposase expression vector (CAG-HyPBase) via a Neon transfection system (Thermo Fisher Scientific, USA) with a 10 μl Neon tip using condition #6 (pulse voltage 1100 V, pulse width 30 ms, pulse no. 1). Then, the cells were plated onto 6-well plates in StemFit medium (AK02N) containing 10 μM Y27632 (Wako, Japan). Two days after transfection, the medium was replaced with fresh medium containing 100 µg/mL G418 (Wako, Japan) or 0.5 µg/mL puromycin (Sigma-Aldrich, USA) and cultured for up to seven days until hiPSC colonies appeared. The medium was changed every other day or every two days. To establish clonal *MYOD1*-hiPSCs (G418- clones, Puro-clones), 12 *MYOD1*-hiPSC colonies were picked from each of the G418- or puromycin-selected plates and expanded in the presence of G418 or puromycin, respectively. Then, 6 *MYOD1*-hiPSC clones that were morphologically better maintained in the undifferentiated state and better proliferated than other clones were selected from each Puro- clone and G418-clone and further expanded for up to two weeks to prepare frozen stocks (Fig. 1B upper panel). To generate bulk *MYOD1*-hiPSCs (G418-bulk line, Puro-bulk line), G418- or puromycin-treated *MYOD1*-hiPSC colonies were bulk-passaged and expanded in the presence of 100 µg/mL G418 or 0.5 µg/mL puromycin, respectively, to prepare frozen stocks (Fig. 1B lower panel). 409B2-*MYOD1*-hiPSCs were established previously via transduction of PB-TA-*hMYOD1*-ERP (22) and were used as control *MYOD1*-hiPSCs. These *MYOD1*- hiPSCs were confirmed to efficiently differentiate into skeletal muscles.

### Differentiation of hiPSCs into skeletal muscles

For differentiation into skeletal muscles, semiconfluent *MYOD1*-hiPSCs maintained under feeder-free conditions were dissociated into single cells by Accutase and plated onto 24-well plates coated with growth factor-reduced Matrigel for 2 hours at a dilution of 1:50 (Corning, USA) at a density of 7 x 10^4^ cells/well in StemFit medium (AK02N) with 10 μM Y27632. On day 1, the medium was changed to hESC medium without FGF-2. On day 3, the medium was replaced with skeletal muscle induction medium consisting of αMEM (Nacalai Tesque, Japan) supplemented with 10% KSR, 2% Ultroser G (BioSepra, Pall, USA), and 0.1 mM 2- ME. To induce the expression of *3HA-hMYOD1*, 1.5 μg/ml Dox (Wako, Japan) was added to the medium from day 2 to day 7. The medium was changed every other day. On day 7, for the formation of mature myotubes, the medium was substituted with myotube differentiation medium consisting of DMEM (high-glucose, 1,500 mg/L NaHCO3, Kohjin Bio, Japan) supplemented with 5% horse serum (HS; Sigma-Aldrich, USA), 10 ng/mL recombinant human insulin-like growth factor 1 (IGF-1) (R&D Systems, USA), and 10 μM SB431542 (Tocris, UK). The medium was changed every two days, and myotube formation was assessed at day 9 of differentiation.

### Culture of C2C12 and Hu5/KD3 cells

C2C12 cells, a mouse myoblast cell line, were cultured as previously described with modifications (44). Briefly, the cells were cultured in DMEM (Nacalai Tesque, Japan) supplemented with 10% fetal bovine serum (FBS) (Sigma-Aldrich, USA) and 1% penicillin-streptomycin (P/S) (Thermo Fisher Scientific, USA) (growth medium). The cells were maintained at 37 °C in a humidified atmosphere containing 5% CO2, and the medium was replaced every day with fresh growth medium. Passaging was conducted when the cells reached 80–90% confluency.

The immortalized human myogenic cell clone Hu5/KD3 was kindly provided by Dr. Naohiro Hashimoto at the National Center for Geriatrics and Gerontology. The Hu5/KD3 cells were maintained and differentiated as previously described with modifications (45–47). Briefly, the cells were cultured in DMEM supplemented with 20% FBS (Nichirei, Japan) and 2% Ultroser G (BioSepra, Pall, USA) at 37 °C in a humidified atmosphere containing 5% CO2. Passaging was conducted when the cells reached 80–90% confluency.

These cell lines were used as controls for qRT-PCR and fabrication of 3D muscle tissues.

### Fabrication of 3D-muscle tissues from *MYOD1*-iPSCs

Fabrication of 3D muscle tissues on microdevices and measurement of contractile force were performed as described previously with modifications (34, 48). Briefly, microdevices made of polydimethylsiloxane (PDMS; SILPOT 184, Dow Corning Toray, Japan) were installed at the center of each well of a 48-well plate, sterilized under UV light for more than 1 hour, and coated with 2% Pluronic F-127 solution (Thermo Fisher Scientific, USA) for 1 hour at room temperature to prevent hydrogel adhesion.

Semiconfluent human *MYOD1*-hiPSCs maintained under feeder-free conditions were dissociated into single cells and differentiated into muscle cells on 6-well plates at a density of 3.5 x 10^5^ cells/well following the same protocol used for monolayer differentiation. From day 2, the expression of the *3HA-hMYOD1 or MYOD1* transgene was induced with 1.5 μg/ml Dox. On day 3, the cells were redissociated by Accutase (22), resuspended in skeletal muscle induction medium, and processed for the fabrication of 3D muscle tissues. A hydrogel mixture was prepared by mixing ice-cold fibrinogen from bovine plasma (10 mg/mL, F8630, Sigma-Aldrich, USA), Matrigel (Corning, USA), and 2× DMEM with a volume ratio of 0.2:0.1:0.2. Then, the cell suspension (6 × 10^6^ cells/ml) and the hydrogel mixture were mixed at a volume ratio of 0.484:0.5. Finally, thrombin from bovine plasma (50 U/ml, Sigma-Aldrich, USA) was added to the mixture at a thrombin:total mixture volume ratio of 0.016:1. Forty-five microliters of the resulting cell and hydrogel solution mixture (1.3 × 10^5^ cells) was poured into the dumbbell-shaped pocket on each microdevice, and the plates were incubated at 37 °C to solidify the hydrogel. The cells applied to the devices were cultured for 4 days in skeletal muscle induction medium with 0.5% P/S in the presence of 1.5 μg/ml Dox (29). The medium was changed every two days. On day 7, the medium was switched to differentiation medium consisting of DMEM, 5% HS, 10 μM SB431542, and 0.5% P/S without Dox, which was changed every other day.

For the fabrication of 3D muscle tissues from C2C12 or Hu5/KD3 cells, semiconfluent myoblast cells were dissociated by using 0.05% trypsin-EDTA, resuspended in growth medium (GM) consisting of DMEM supplemented with 10% FBS, and processed as reported previously (48). The cell suspension (2 × 10^6^ cells/ml) and the hydrogel mixture were mixed following the same protocol used for the hiPSCs. The tissues were cultured in GM for two days, and the medium was then switched to differentiation medium consisting of DMEM supplemented with 2% HS, 1% P/S, and 1% ITS (Sigma-Aldrich, USA). On day 6, muscle tissues were pulled up at the top of the pillar and processed for the measurement of contractile force after EPS.

All the culture media contained 2.0 mg/mL 6-aminocaproic acid (Sigma-Aldrich, USA) and 1 mg/ml trans-4-aminomethyl cyclohexane carboxylic acid (TA) (Tokyo Chemical Industries, Japan) to prevent disassembly of the tissues.

### Contractile force measurement of 3D muscle tissues

The contractile force of the 3D muscle tissues was measured as reported previously (34). Muscle tissues were electrically stimulated to achieve maximum tetanic force with an electrical stimulus of 4.0 V/mm at 30 Hz with 2 msec wide pulses (C-Pace EP, IonOptix, USA) using customized electrodes. Electrical stimulation was applied twice to each tissue sample for 5 seconds on days 11, 13, 15, and 17, and the displacement of the tips of the micro-posts was observed with an upright microscope (BX53F, Olympus, Japan) under a 4x objective (UPLFN4X) or a 20x objective (SLMPLN20X). The displacement of the tip of each micro-post was measured with ImageJ software using photographs taken with and without stimulation, and the average value from both micro-posts was obtained. The contractile force (F) was determined by the equation F = 3πER^4^δ/(4L^3^), where E is the elastic modulus of PDMS (1.7 MN/m^2^), R is the radius of a micro-post (0.25 mm), δ is the displacement of the tip of the micro-post, and L is the length of the micro-post (4 mm) (49).

### Histological analysis

ICC and IHC analyses of cultured cells and 3D muscle tissues, respectively, were performed as described previously with modifications (34, 50). Cultured cells were fixed with 4% paraformaldehyde (PFA) for 15 min at room temperature. After blocking in blocking buffer consisting of phosphate-buffered saline (PBS) containing 10% FBS and 0.3% Triton X-100 for 1 hour at room temperature, the cells were incubated with primary antibodies at 4 °C overnight (see Supplementary Table S2 for the primary antibodies). After three washes with PBS, the cells were incubated with secondary antibodies conjugated with Alexa 488, Alexa 555, or Alexa 647 for 1 hour at room temperature. For the quantitative analysis, photographs of five randomly selected visual fields were taken and used for quantification of the immunopositive cells and nuclei and measurement of the immune-positive area and myotube thickness manually or with ImageJ software (51).

Muscle tissues were fixed with 4% PFA for 30 min at room temperature. After permeabilization with PBS containing 0.3% Triton X-100 for 10 min, the tissues were incubated in blocking buffer consisting of PBS containing 10% goat serum and 0.01% Triton X-100 for 30 min at room temperature and then incubated with primary antibodies (see Supplementary Table S2 for the primary antibodies) for 3 hours at room temperature. After three washes with PBS, the cells were incubated with secondary antibodies conjugated with Alexa 488 or Alexa 555 for 2 hours at room temperature.

Both the cells and the muscle tissues were stained for nuclei with 10 μg/ml Hoechst 33258 (Sigma-Aldrich, USA) and were observed with a confocal laser scanning microscope (LSM700 or LSM900, Carl Zeiss, Germany).

### RNA isolation and qRT-PCR analysis

RNA was isolated using an RNAeasy Mini Kit (Qiagen, Germany) and then converted into cDNA using PrimeScript RT Master Mix (Takara Bio Inc, Japan). Real-time qRT-PCR was performed as previously described using TB Green Premix Ex Taq II (Takara Bio Inc, Japan) and a QuantStudio 7 Real-Time PCR system (Thermo Fisher Scientific, USA) (52). The amount of cDNA was normalized to that of human-specific *β-ACTIN* mRNA. The primer sequences and PCR cycling conditions are listed in Supplementary Table S1.

### Statistical analysis

The data are presented as the mean ± standard error of the mean (SEM). Statistical analyses were performed by one-way analysis of variance (ANOVA) followed by a *post hoc* Bonferroni test. A *p* value less than 0.05 was considered to indicate statistical significance.

## Study approval

All the experimental procedures involving the use of hiPSCs were approved by the ethics committee of the Aichi Medical University School of Medicine (approval numbers 2020- H072 and 2020-413).

## Author Contributions

MIR, TI, and YO conceived and designed the study. MIR, TI, DS, and YO performed the experiments and analyzed the data. HS provided 409B2-*MYOD1*-hiPSCs. HO provided EKN3-hiPSCs. KA, TN, KY, and KS supported the fabrication of 3D muscle tissues and the measurement of contractile force. KO, RO, and ZK provided technical assistance. MIR and YO wrote the manuscript. HO, HS, KS and MD provided technical assistance, scientific discussion, and critical reading. All authors read and approved the final manuscript.

## Acknowledgments

We are grateful to Dr. S. Yamanaka (Kyoto University) for hiPSCs (201B7, 409B2); to Dr. K. Woltjen (Kyoto University) for the *piggyBac* vector for Dox-inducible expression; to Dr. N. Hashimoto (National Center for Geriatrics and Gerontology) for the human myoblast cell line Hu5/KD3; to the Aichi Medical University Institute of Comprehensive Medical Research Division of Advanced Research Promotion for the assistance in DNA sequence analyses; to Ms. M. Ishihara, Ms. E. Mitani, and Ms. A. Hashimoto for technical and administrative support; and to all members of Dr. Okada’s laboratory for encouragement and support. This work was supported by the Japan Society for the Promotion of Science (JSPS) KAKENHI grant numbers JP17K19465, JP18K15470, JP18K07539, JP19H03576, JP19K07969, JP19J40240, JP20K16611, and JP21K07471 (to TI, MD, and YO); by the Japan Agency for Medical Research and Development (AMED) under grant numbers JP19ek0109243 and JP21bm0804020 (to TI, MD, and YO); by a grant from the Japan SBMA Association (to TI and KO); and by a grant from the Hori Sciences and Arts Foundation (to TI). MIR and ZK are students sponsored by the Japanese government (Ministry of Education, Culture, Sports, Science and Technology; MEXT).

## References

1. Takahashi K, and Yamanaka S. Induction of pluripotent stem cells from mouse embryonic and adult fibroblast cultures by defined factors. Cell. 2006;126(4):663–76.

2. Takahashi K, Tanabe K, Ohnuki M, Narita M, Ichisaka T, Tomoda K, et al. Induction of pluripotent stem cells from adult human fibroblasts by defined factors. Cell. 2007;131(5):861–72.

3. Corti S, Faravelli I, Cardano M, and Conti L. Human pluripotent stem cells as tools for neurodegenerative and neurodevelopmental disease modeling and drug discovery. Expert Opin Drug Discov. 2015;10(6):615–29.

4. Chal J, and Pourquié O. Making muscle: skeletal myogenesis in vivo and in vitro. Development. 2017;144(12):2104–22.

5. Xu C, Tabebordbar M, Iovino S, Ciarlo C, Liu J, Castiglioni A, et al. A zebrafish embryo culture system defines factors that promote vertebrate myogenesis across species. Cell. 2013;155(4):909–21.

6. Hosoyama T, McGivern JV, Van Dyke JM, Ebert AD, and Suzuki M. Derivation of myogenic progenitors directly from human pluripotent stem cells using a sphere-based culture. Stem Cells Transl Med. 2014;3(5):564–74.

7. Borchin B, Chen J, and Barberi T. Derivation and FACS-mediated purification of PAX3+/PAX7+ skeletal muscle precursors from human pluripotent stem cells. Stem Cell Reports. 2013;1(6):620–31.

8. Choi IY, Lim H, Estrellas K, Mula J, Cohen TV, Zhang Y, et al. Concordant but Varied Phenotypes among Duchenne Muscular Dystrophy Patient-Specific Myoblasts Derived using a Human iPSC-Based Model. Cell Rep. 2016;15(10):2301–12.

9. Chal J, Al Tanoury Z, Hestin M, Gobert B, Aivio S, Hick A, et al. Generation of human muscle fibers and satellite-like cells from human pluripotent stem cells in vitro. Nat Protoc. 2016;11(10):1833–50.

10. van der Wal E, Herrero-Hernandez P, Wan R, Broeders M, In ’t Groen SLM, van Gestel TJM, et al. Large-Scale Expansion of Human iPSC-Derived Skeletal Muscle Cells for Disease Modeling and Cell-Based Therapeutic Strategies. Stem Cell Reports. 2018;10(6):1975–90.

11. Davis RL, Weintraub H, and Lassar AB. Expression of a single transfected cDNA converts fibroblasts to myoblasts. Cell. 1987;51(6):987–1000.

12. Warren L, Manos PD, Ahfeldt T, Loh YH, Li H, Lau F, et al. Highly efficient reprogramming to pluripotency and directed differentiation of human cells with synthetic modified mRNA. Cell Stem Cell. 2010;7(5):618–30.

13. Darabi R, Arpke RW, Irion S, Dimos JT, Grskovic M, Kyba M, et al. Human ES- and iPS-derived myogenic progenitors restore DYSTROPHIN and improve contractility upon transplantation in dystrophic mice. Cell Stem Cell. 2012;10(5):610–9.

14. Goudenege S, Lebel C, Huot NB, Dufour C, Fujii I, Gekas J, et al. Myoblasts derived from normal hESCs and dystrophic hiPSCs efficiently fuse with existing muscle fibers following transplantation. Mol Ther. 2012;20(11):2153–67.

15. Rao L, Tang W, Wei Y, Bao L, Chen J, Chen H, et al. Highly efficient derivation of skeletal myotubes from human embryonic stem cells. Stem Cell Rev Rep. 2012;8(4):1109–19.

16. Tanaka A, Woltjen K, Miyake K, Hotta A, Ikeya M, Yamamoto T, et al. Efficient and reproducible myogenic differentiation from human iPS cells: prospects for modeling Miyoshi Myopathy in vitro. PLoS One. 2013;8(4):e61540.

17. 17. Tedesco FS, Gerli MF, Perani L, Benedetti S, Ungaro F, Cassano M, et al. Transplantation of genetically corrected human iPSC-derived progenitors in mice with limb-girdle muscular dystrophy. Sci Transl Med. 2012;4(140):140ra89.

18. Lenzi J, Pagani F, De Santis R, Limatola C, Bozzoni I, Di Angelantonio S, et al. Differentiation of control and ALS mutant human iPSCs into functional skeletal muscle cells, a tool for the study of neuromuscolar diseases. Stem Cell Res. 2016;17(1):140–7.

19. Shoji E, Woltjen K, and Sakurai H. Directed Myogenic Differentiation of Human Induced Pluripotent Stem Cells. Methods Mol Biol. 2016;1353:89–99.

20. Maffioletti SM, Gerli MF, Ragazzi M, Dastidar S, Benedetti S, Loperfido M, et al. Efficient derivation and inducible differentiation of expandable skeletal myogenic cells from human ES and patient-specific iPS cells. Nat Protoc. 2015;10(7):941–58.

21. Filareto A, Parker S, Darabi R, Borges L, Iacovino M, Schaaf T, et al. An ex vivo gene therapy approach to treat muscular dystrophy using inducible pluripotent stem cells. Nat Commun. 2013;4:1549.

22. Uchimura T, Otomo J, Sato M, and Sakurai H. A human iPS cell myogenic differentiation system permitting high-throughput drug screening. Stem Cell Res. 2017;25:98–106.

23. Yoshida T, Awaya T, Jonouchi T, Kimura R, Kimura S, Era T, et al. A Skeletal Muscle Model of Infantile-onset Pompe Disease with Patient-specific iPS Cells. Sci Rep. 2017;7(1):13473.

24. Rao L, Qian Y, Khodabukus A, Ribar T, and Bursac N. Engineering human pluripotent stem cells into a functional skeletal muscle tissue. Nat Commun. 2018;9(1):126.

25. 25. Abujarour R, Bennett M, Valamehr B, Lee TT, Robinson M, Robbins D, et al. Myogenic differentiation of muscular dystrophy-specific induced pluripotent stem cells for use in drug discovery. Stem Cells Transl Med. 2014;3(2):149–60.

26. Albini S, Coutinho P, Malecova B, Giordani L, Savchenko A, Forcales SV, et al. Epigenetic reprogramming of human embryonic stem cells into skeletal muscle cells and generation of contractile myospheres. Cell Rep. 2013;3(3):661–70.

27. Skoglund G, Lainé J, Darabi R, Fournier E, Perlingeiro R, and Tabti N. Physiological and ultrastructural features of human induced pluripotent and embryonic stem cell-derived skeletal myocytes in vitro. Proc Natl Acad Sci U S A. 2014;111(22):8275–80.

28. Buckingham M, and Rigby PW. Gene regulatory networks and transcriptional mechanisms that control myogenesis. Dev Cell. 2014;28(3):225–38.

29. Berkes CA, and Tapscott SJ. MyoD and the transcriptional control of myogenesis. Semin Cell Dev Biol. 2005;16(4-5):585–95.

30. Shimojo D, Onodera K, Doi-Torii Y, Ishihara Y, Hattori C, Miwa Y, et al. Rapid, efficient, and simple motor neuron differentiation from human pluripotent stem cells. Mol Brain. 2015;8(1):79.

31. Matsumoto T, Fujimori K, Andoh-Noda T, Ando T, Kuzumaki N, Toyoshima M, et al. Functional Neurons Generated from T Cell-Derived Induced Pluripotent Stem Cells for Neurological Disease Modeling. Stem Cell Reports. 2016;6(3):422–35.

32. Onodera K, Shimojo D, Ishihara Y, Yano M, Miya F, Banno H, et al. Unveiling synapse pathology in spinal bulbar muscular atrophy by genome-wide transcriptome analysis of purified motor neurons derived from disease specific iPSCs. Mol Brain. 2020;13(1):18.

33. Swist S, Unger A, Li Y, Vöge A, von Frieling-Salewsky M, Skärlén Å, et al. Maintenance of sarcomeric integrity in adult muscle cells crucially depends on Z-disc anchored titin. Nat Commun. 2020;11(1):4479.

34. Nagashima T, Hadiwidjaja S, Ohsumi S, Murata A, Hisada T, Kato R, et al. In Vitro Model of Human Skeletal Muscle Tissues with Contractility Fabricated by Immortalized Human Myogenic Cells. Adv Biosyst. 2020;4(11):e2000121.

35. Osaki T, Uzel SGM, and Kamm RD. Microphysiological 3D model of amyotrophic lateral sclerosis (ALS) from human iPS-derived muscle cells and optogenetic motor neurons. Sci Adv. 2018;4(10):eaat5847.

36. Maffioletti SM, Sarcar S, Henderson ABH, Mannhardt I, Pinton L, Moyle LA, et al. Three-Dimensional Human iPSC-Derived Artificial Skeletal Muscles Model Muscular Dystrophies and Enable Multilineage Tissue Engineering. Cell Rep. 2018;23(3):899–908.

37. Vara JA, Portela A, Ortín J, and Jiménez A. Expression in mammalian cells of a gene from Streptomyces alboniger conferring puromycin resistance. Nucleic Acids Res. 1986;14(11):4617–24.

38. Southern PJ, and Berg P. Transformation of mammalian cells to antibiotic resistance with a bacterial gene under control of the SV40 early region promoter. J Mol Appl Genet. 1982;1(4):327–41.

39. Watanabe S, Kai N, Yasuda M, Kohmura N, Sanbo M, Mishina M, et al. Stable production of mutant mice from double gene converted ES cells with puromycin and neomycin. Biochem Biophys Res Commun. 1995;213(1):130–7.

40. Guo C, Fordjour FK, Tsai SJ, Morrell JC, and Gould SJ. Choice of selectable marker affects recombinant protein expression in cells and exosomes. J Biol Chem. 2021;297(1):100838.

41. Fujimori K, Ishikawa M, Otomo A, Atsuta N, Nakamura R, Akiyama T, et al. Modeling sporadic ALS in iPSC-derived motor neurons identifies a potential therapeutic agent. Nat Med. 2018;24(10):1579–89.

42. Kim SI, Oceguera-Yanez F, Sakurai C, Nakagawa M, Yamanaka S, and Woltjen K. Inducible Transgene Expression in Human iPS Cells Using Versatile All-in-One piggyBac Transposons. Methods Mol Biol. 2016;1357:111–31.

43. Okita K, Matsumura Y, Sato Y, Okada A, Morizane A, Okamoto S, et al. A more efficient method to generate integration-free human iPS cells. Nat Methods. 2011;8(5):409–12.

44. Yaffe D, and Saxel O. Serial passaging and differentiation of myogenic cells isolated from dystrophic mouse muscle. Nature. 1977;270(5639):725-7.

45. Wada MR, Inagawa-Ogashiwa M, Shimizu S, Yasumoto S, and Hashimoto N. Generation of different fates from multipotent muscle stem cells. Development. 2002;129(12):2987–95.

46. Hashimoto N, Kiyono T, Wada MR, Shimizu S, Yasumoto S, and Inagawa M. Immortalization of human myogenic progenitor cell clone retaining multipotentiality. Biochem Biophys Res Commun. 2006;348(4):1383–8.

47. Shiomi K, Kiyono T, Okamura K, Uezumi M, Goto Y, Yasumoto S, et al. CDK4 and cyclin D1 allow human myogenic cells to recapture growth property without compromising differentiation potential. Gene Ther. 2011;18(9):857–66.

48. Shimizu K, Genma R, Gotou Y, Nagasaka S, and Honda H. Three-Dimensional Culture Model of Skeletal Muscle Tissue with Atrophy Induced by Dexamethasone. Bioengineering (Basel). 2017;4(2).

49. Vandenburgh H, Shansky J, Benesch-Lee F, Barbata V, Reid J, Thorrez L, et al. Drug-screening platform based on the contractility of tissue-engineered muscle. Muscle Nerve. 2008;37(4):438–47.

50. Okada Y, Matsumoto A, Shimazaki T, Enoki R, Koizumi A, Ishii S, et al. Spatiotemporal recapitulation of central nervous system development by murine embryonic stem cell-derived neural stem/progenitor cells. Stem Cells. 2008;26(12):3086–98.

51. Schneider CA, Rasband WS, and Eliceiri KW. NIH Image to ImageJ: 25 years of image analysis. Nat Methods. 2012;9(7):671–5.

52. Okada R, Onodera K, Ito T, Doyu M, Okano HJ, and Okada Y. Modulation of oxygen tension, acidosis, and cell density is crucial for neural differentiation of human induced pluripotent stem cells. Neurosci Res. 2021;163:34–42.

