## Supplemental Materials for "Simple and efficient differentiation of human iPSCs into contractible skeletal muscles for muscular disease modeling"

**Supplementary Table S1. Primer sequences and cycling conditions for qRT-PCR**

| *Genes* | Primer name | Primer sequence | Temp. | Product size |
|---|---|---|---|---|
| *OCT3/4* | hPOU5F1-S767 | AGAACCGAGTGAGAGGCAAC | 62 | 206 |
| *(hPOU5F1)* | hPOU5F1-AS972 | TGAGAAAGGAGACCCAGCAG | | |
| *NANOG* | hNANOG-S286 | CCTATGCCTGTGATTTGTGG | 62 | 218 |
| | hNANOG-AS503 | TGTTTCTTGACTGGGACCTTG | | |
| *CD56* | hNCAM1-S351 | CAGCCAGCAGATTACAATGC | 62 | 99 |
| | hNCAM1-AS449 | TGGCTGGGAACAATATCCAC | | |
| *MEF2C* | hMEF2C-S3999 | CCACAATTAGACCACAATGCACC | 62 | 137 |
| | hMEF2C-AS4135 | TCCATGGATACCTTGAGTAGTGG | | |
| *TMEM8C* | hTMEM8C-S433 | CCAGACAAGAGCGTCTACACC | 62 | 81 |
| | hTMEM8C-AS513 | AAAGAAGAAGCGTAGCATCAGG | | |
| *Tg MYOD1* | hMYOD1-S815 | CGGACGTGCCTTCTGAGTC | 62 | 184 |
| | Tg-hMYOD-AS | CTTTGTACAAGAAAGCTGGGTC | | |
| *Total MYOD1* | hMYOD1-S815 | CGGACGTGCCTTCTGAGTC | 62 | 145 |
| | hMYOD1-AS959 | AGCACCTGGTATATCGGGTTG | | |
| *Endo MYOD1* | hMYOD-S1331、 | CACTCCGGTCCCAAATGTAG | 62 | 180 |
| | hMYOD- AS1511 | TTCCCTGTAGCACCACAGAG | | |
| *MYF6* | hMYF6-S612 | AGGAGCAAGTATTGATTCGTCAG | 62 | 89 |
| | hMYF6-AS700 | GGAGTTTGCGTTCCTCCGAG | | |
| *MYH2* | hMYH2-S68 | CTGTCTCACTCCCAGGCTAC | 62 | 159 |
| | hMYH2-AS226 | TCAAAGGGCCTATTCTGGGC | | |
| *MYH7* | hMYH7-S5725 | AAGGTCAAGGCCTACAAGCG | 62 | 178 |
| | hMYH7-AS5902 | TCAAGCCCTTCGTGCCAATG | | |
| *β-ACTIN* | hβ-ACTIN-S1062 | GATCAAGATCATTGCTCCTCCT | 62 | 180 |
| | hβ-ACTIN-AS1241 | GGGTGTAACGCAACTAAGTCA | | |
| *MYOG* | hMYOG-S378 | TTCGAGGCCCTGAAGAGAAG | 62 | 292 |
| | hMYOG-AS669 | GGTTGTGGGCATCTGTAGGG | | |
| *MYF5* | hMYF5-S128 | TCAGCAGGATGGACGTGATG | 62 | 102 |
| | hMYF5-AS229 | ACTCGTCCCCAAATTCACCC | | |

Temp: Annealing temperature

**Supplementary Table S2. Antibodies used in this study**

| Antigen | Host | Isotype | Dilution | Cat. no / Clone Name | Manufacturer |
|---|---|---|---|---|---|
| MyoD | Ms | IgG1 | 1:500 | 5.8A | DAKO |
| HA-Tag | Ms | IgG2b | 1:1000 | TANA2 | MBL |
| MHC | Ms | IgG2b | 1:500 | MF20 | DSHB |
| MyoG | Ms | IgG2a | 1:500 | F5D | DSHB |
| OCT3/4 | Ms | IgG | 1:100 | SC-5279 | Santa Cruz |
| Nanog | Rb | IgG | 1:100 | RCAB003P | ReproCELL |
| Titin | Ms | IgM | 1:500 | 9D10 | DSHB |
| $\alpha$-Actinin | Ms | IgG1 | 1:500 | A7811 | Sigma-Aldrich |

*DSHB: Developmental Studies Hybridoma Bank

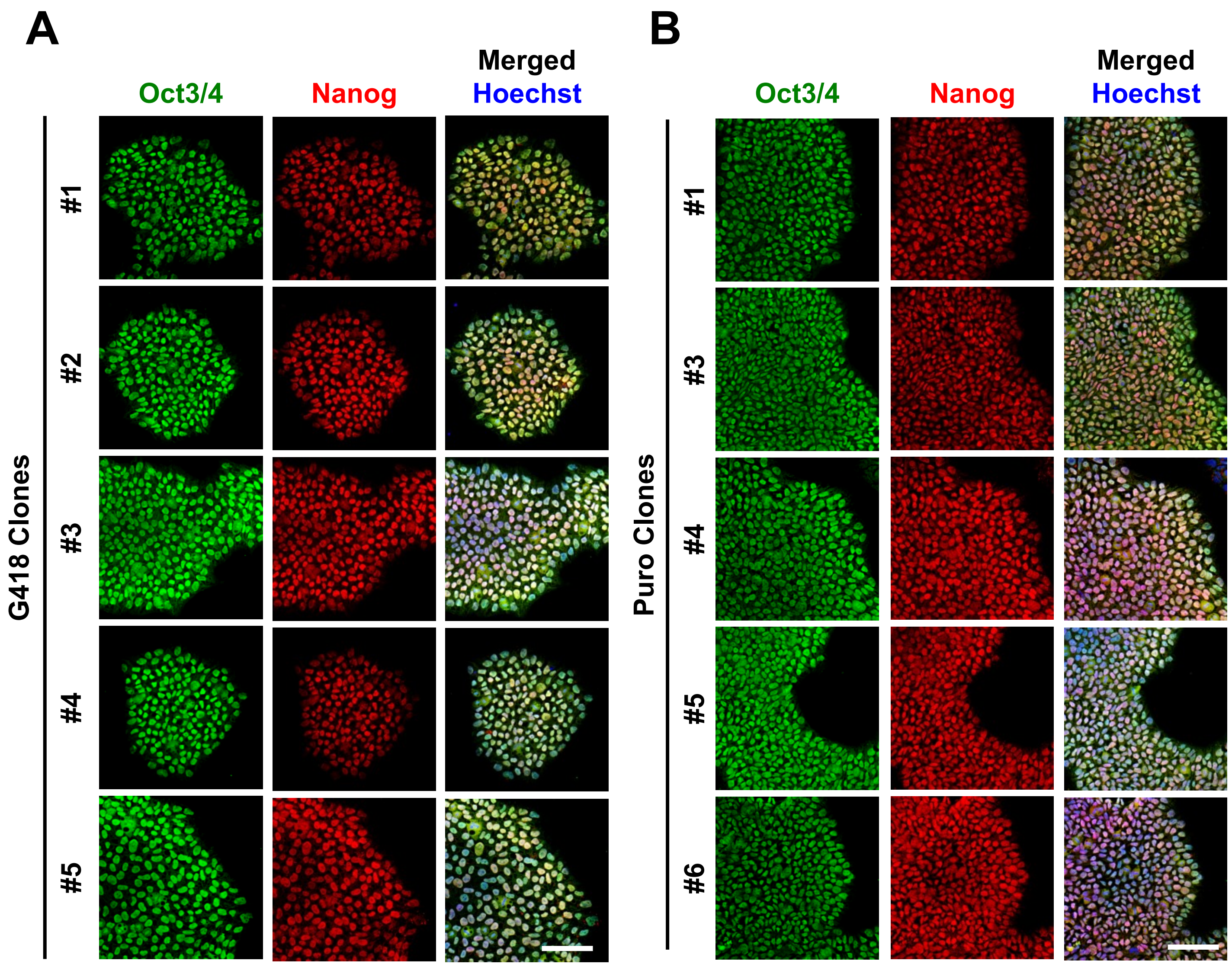


**Supplementary Figure S1 The expression of pluripotent stem cell markers in *MYOD1*-hiPSCs.**
A, B. ICC analysis of the clonal *MYOD1*-hiPSCs established with G418 (A) or puromycin selection (B) for pluripotent stem cell markers (Oct3/4 and Nanog). The nuclei were stained with Hoechst 33258. All *MYOD1*-hiPSCs retained the expression of pluripotent stem cell markers. Scale bar, 50 μm.

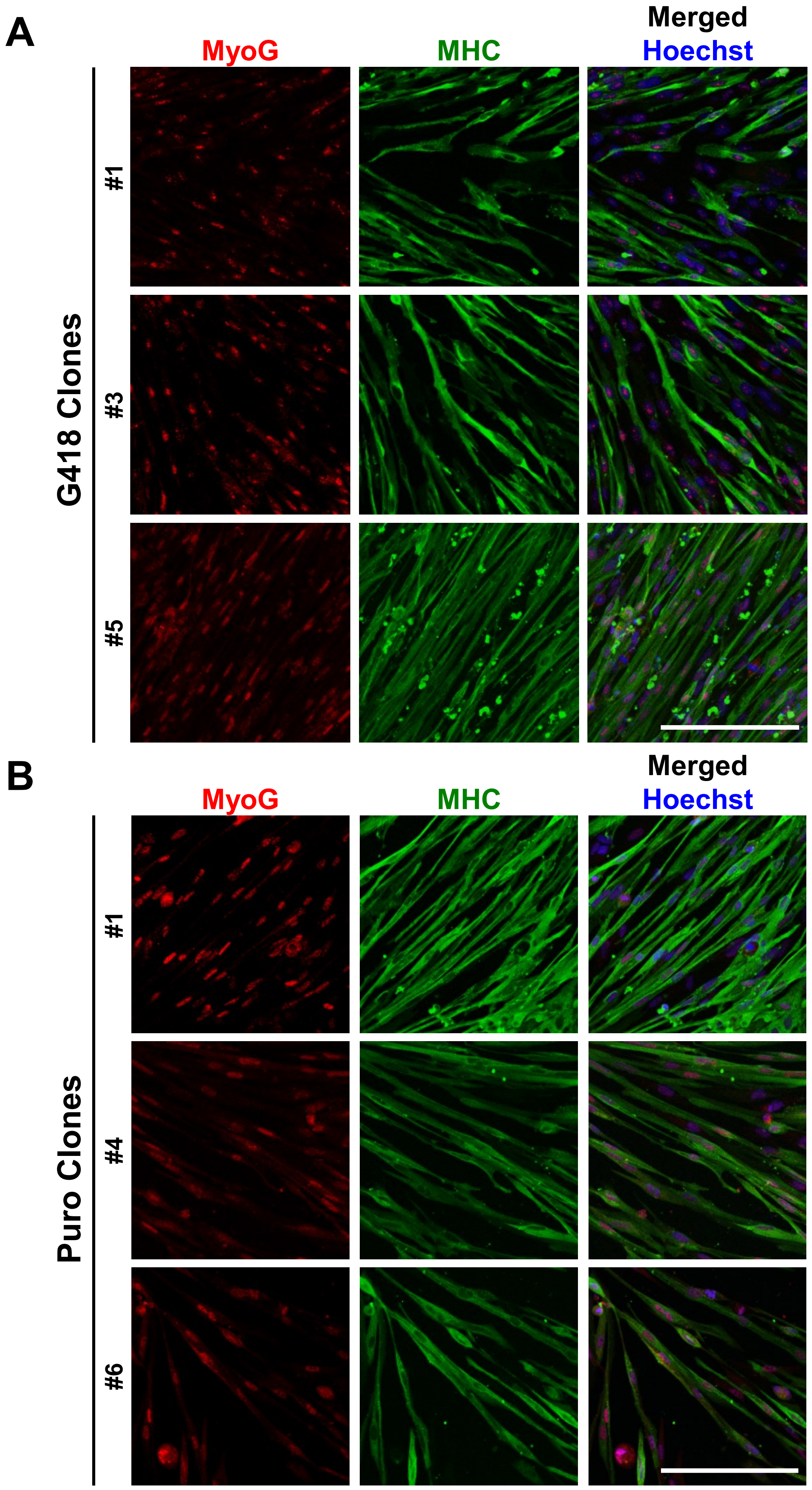


**Supplementary Figure S2 Differentiation potentials of clonal *MYOD1*-hiPSCs established by G418 or puromycin.**

A, B. ICC analysis of myotubes derived from clonal *MYOD1*-hiPSCs established by G418 or puromycin selection for the expression of MyoG and MHC at day 9 of differentiation. The nuclei were stained with Hoechst 33258. Scale bar, 200 µm.

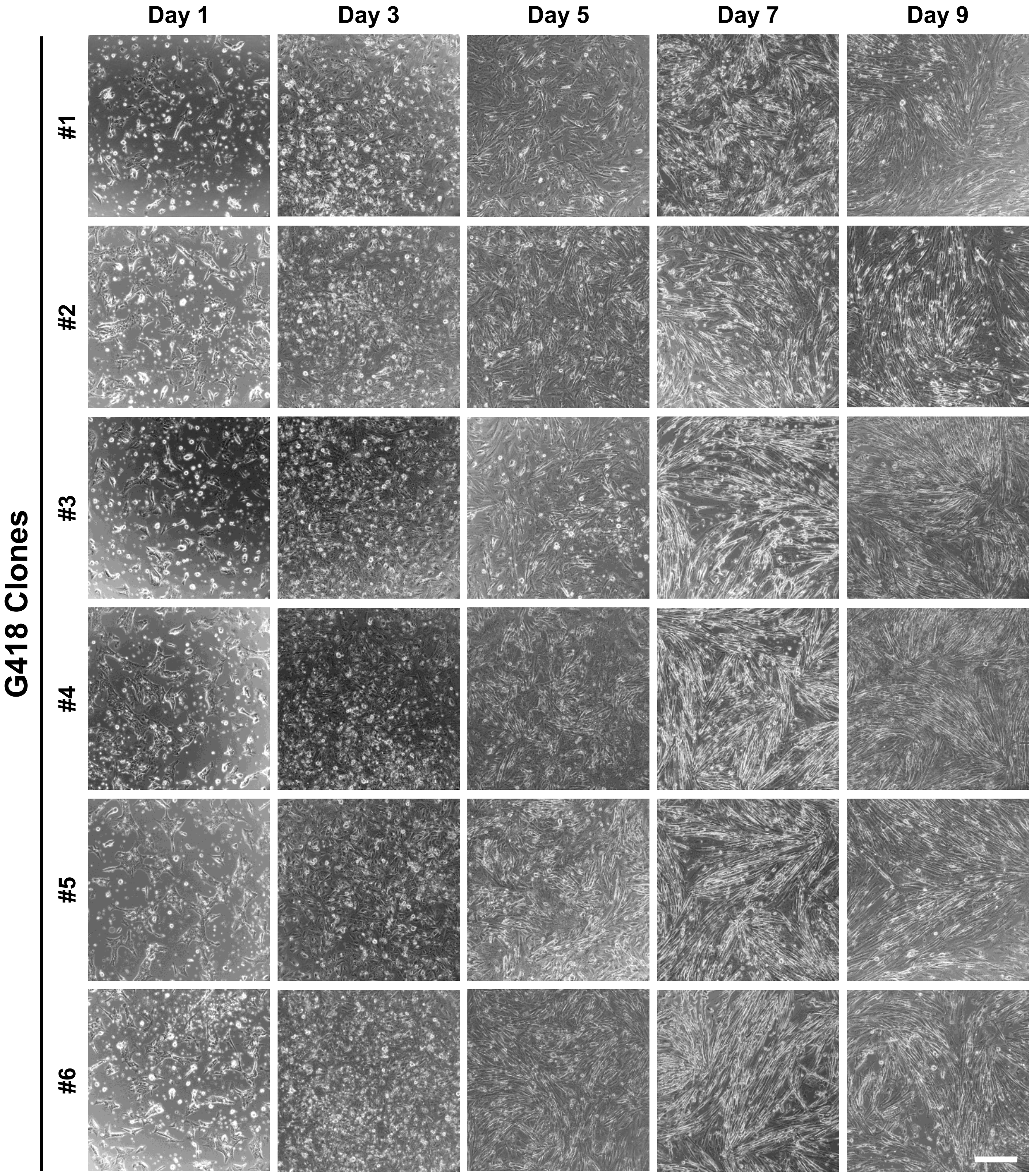


**Supplementary Figure S3 Time course muscular differentiation of clonal *MYOD1*-hiPSCs established with G418 selection.**

Brightfield images showing the time course of muscular differentiation of the established clonal *MYOD1*-hiPSCs with G418 selection. All clonal *MYOD1*-hiPSCs morphologically differentiated into myoblasts, subsequently formed myotubes, and achieved a myotube-like aligned structure by day 7. Scale bar, 100 µm.

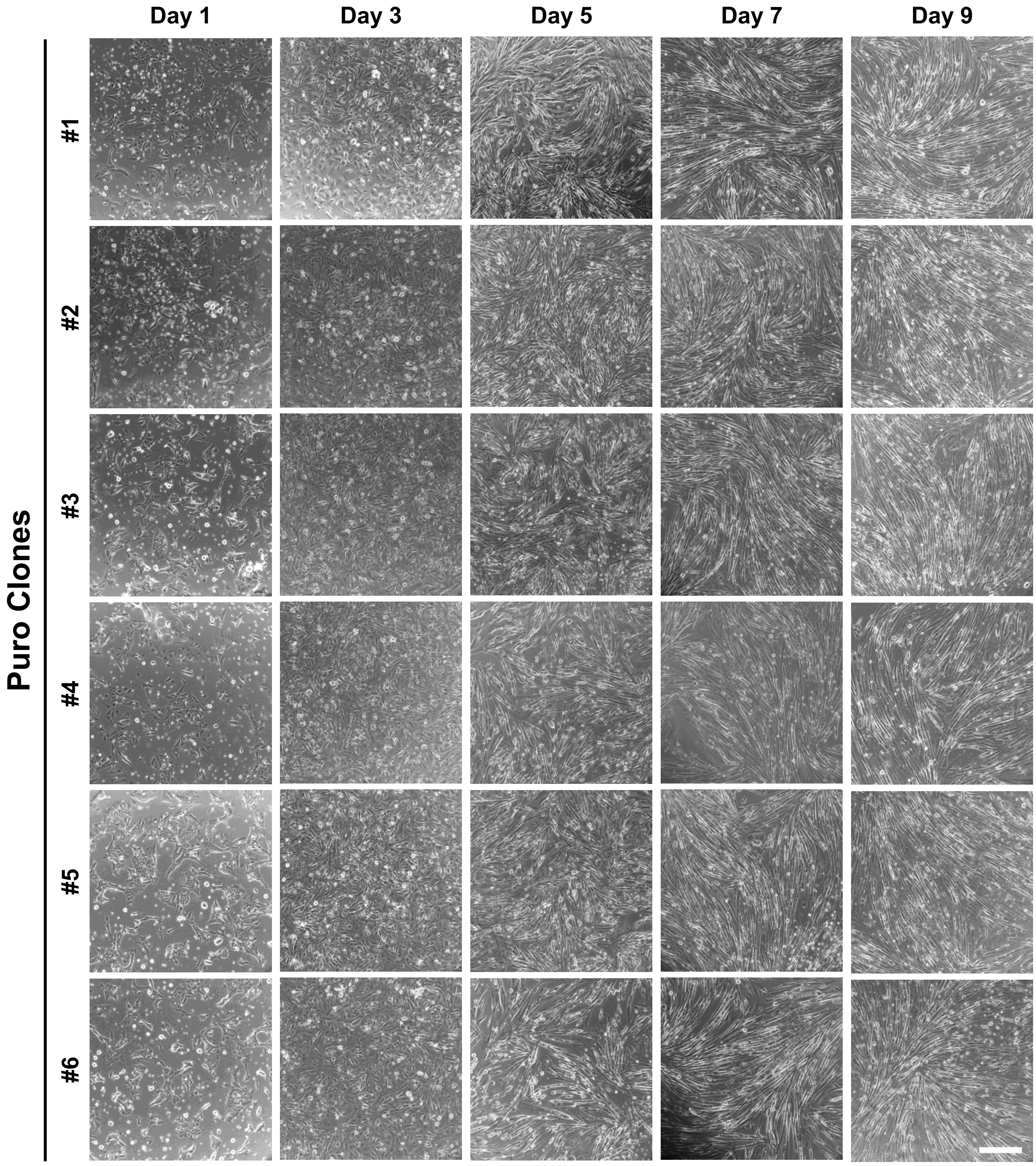


**Supplementary Figure S4 Time course muscular differentiation of clonal *MYOD1*-hiPSCs established with puromycin selection.**

Brightfield images showing the time course of muscular differentiation of the established clonal *MYOD1*-hiPSCs with puromycin selection. All clonal *MYOD1*-hiPSCs morphologically differentiated into myoblasts, subsequently formed myotubes, and achieved a myotube-like aligned structure by day 7. Scale bar, 100 µm.

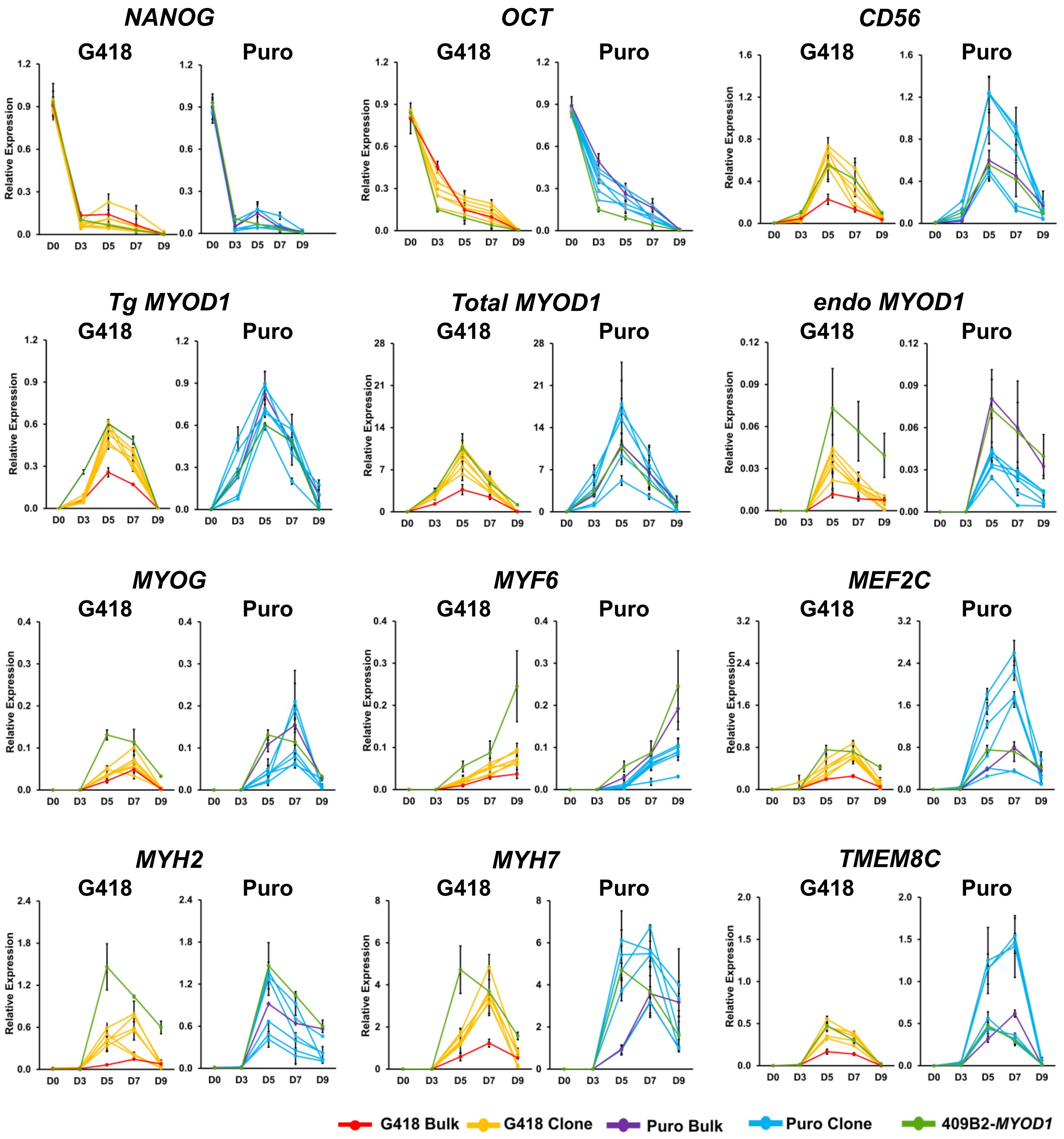


**Supplementary Figure S5 Time course gene expression of bulk and clonal *MYOD1*-hiPSCs with G418 or puromycin selection.**

Time course gene expression analysis of bulk and clonal *MYOD1*-hiPSCs with G418 or puromycin selection as well as control 409B2-*MYOD1*-hiPSCs along with muscular differentiation. Compared with the Puro-clones and 409B2-*MYOD1*-hiPSCs, the Puro-bulk line showed similar or average expression, whereas the G418-bulk line showed lower expression levels than other *MYOD1*-hiPSCs throughout the differentiation process. Moreover, in contrast to the large clonal variations observed in differentiating Puro-clones, the Puro-bulk line showed average expression of the six Puro-clones or expression similar to that in the G418-clones for all the genes associated with skeletal muscle differentiation during the differentiation processes. The amount of cDNA was normalized to that of human-specific *β-ACTIN* and is presented as the relative expression of undifferentiated hiPSCs (*NANOG* and *OCT3/4*), the human myoblast cell line Hu5/KD3 differentiated for 3 days (*CD56*, total and endogenous *MYOD1*, *MYOG*, *MYF6*, *MEF2C*, *MYH2*, *MYH7*, and *TMEM8C*), and EKN3-*MYOD1* iPSCs differentiated for 5 days (*Tg MYOD1*). The data are presented as the mean ± SEM, n = 3. *, $p < 0.05$, **, $p < 0.01$.
